# Single-molecule imaging and molecular dynamics simulations reveal early activation of the MET receptor *in situ*

**DOI:** 10.1101/2023.12.22.572978

**Authors:** Yunqing Li, Serena Arghittu, Marina S. Dietz, Gabriel J. Hella, Daniel Haße, Davide M. Ferraris, Petra Freund, Hans-Dieter Barth, Hartmut H. Niemann, Roberto Covino, Mike Heilemann

**Author notes:** These authors equally contributed to this work (Yunqing Li, Serena Arghittu).

## Abstract

The assembly of membrane receptors into signaling complexes is at the origin of key cellular events. Yet, we often lack detailed structural mechanistic understanding. Receptors are embedded into a complex cellular membrane, which defines their dynamics but also complicates their experimental characterizations significantly. Here, we showcase an integrative structural biology approach to investigate the activation mechanism of the human growth factor receptor MET. MET is a receptor tyrosine kinase involved in cell proliferation, migration, and survival. MET is also hijacked by the intracellular pathogen *Listeria monocytogenes*. Its invasion protein, internalin B (InlB), binds to MET and promotes the formation of a signaling dimer that triggers the internalization of the pathogen. Crystallography had suggested two different 2:2 MET:InlB complexes. Here, we use a combination of structural biology, modeling, molecular dynamics simulations, and *in situ* single-molecule Förster resonance energy transfer (smFRET) to elucidate the early events in MET activation. Simulations show that InlB binding stabilizes MET in a conformation that promotes dimer formation. smFRET identifies the organization of the *in situ* signaling dimer, which resembles one of the two crystal structures yet shows differences. Further MD simulations resulted in a refinement of the dimer model, which is in quantitative agreement with smFRET results. We accurately describe the structural dynamics underpinning an important cellular event and introduce a powerful methodological pipeline applicable to studying the activation of other plasma membrane receptors *in situ*.

## Introduction

The formation of oligomers of single-pass transmembrane (TM) receptors activates numerous fundamental cellular events. Oligomerization is often induced by ligand-binding, and its stabilization involves weak interactions along the whole length of the receptor-ligand complex, including the receptor-bound ligand, the extracellular domain (ECD), the TM domain, as well as intracellular regions such as the juxtamembrane (JM) or kinase domain ^1–4^. The membrane anchorage of receptors impacts their dynamics significantly. It increases receptors’ local concentration, reduces translational and rotational freedom, and pre-orients the receptor sswcorrectly for oligomerization. Other biomolecules defining the native membrane environment can also contribute to the correct assembly of a receptor-ligand complex, e.g., specific lipids or glycosaminoglycans. To fully understand how a receptor forms biologically active oligomers therefore requires studying the receptor in its native, complex membrane environment. However, this severely complicates a structural and biochemical analysis.

*In vitro* approaches that study soluble fragments, like the ligand-bound receptor ECD, while very informative may not be able to capture the contribution that weak interactions have to complex formation of membrane-embedded proteins. On the other hand, only few methods allow *in situ* analysis of membrane proteins in a native membrane environment. While cryo-electron tomography (cryo-ET) holds great promise for the future ^5^, fluorescence microscopy has already provided important insights ^6–9^. Cutting-edge microscopy methods that can address single proteins in the context of an intact cell ideally complement *in vitro* methods, especially if structural models exist that allow strategic site-specific fluorophore labeling. However, these methods usually lack the structural resolutions necessary to formulate precise mechanistic hypotheses.

Here, we overcome this challenge by introducing an integrative structural biology approach combining computational structural modeling, molecular dynamics (MD) simulations, and single-molecule microscopy experiments to reveal the structural dynamics of membrane receptors *in situ*. We applied this approach to the receptor tyrosine kinase MET, which functions as a signaling protein on the plasma membrane and regulates cell proliferation, migration, and wound healing ^10,11^. Dysfunction of MET is observed for a variety of diseases, such as cancer ^12^, diabetes ^13^ and autism ^14^. In addition, the bacterial pathogen *Listeria monocytogenes* targets MET to initiate host cell invasion ^15^. Signaling of MET is initiated by binding of the physiological ligand HGF ^16,17^, its natural isoform NK1 ^17,18^ or the bacterial ligand InlB. Upon ligand binding, two MET receptors and two ligands assemble into a 2:2 complex, which facilitates trans-phosphorylation of the two MET proteins within the complex and downstream signaling ^19^. However, for both the endogenous and the bacterial ligand, the *in situ* structures and the structural dynamics of the activation mechanism have not been resolved yet.

Using our integrative structural biology approach, we investigated the mechanism of InlB-mediated MET activation *in situ*. Simulations showed that binding of InlB induces an extended conformation of the MET stalk that facilitates the formation of a complex mediated by ligand-ligand contacts. Single-molecule FRET in U-2 OS cells revealed the organization of the 2:2 (MET:InlB)_2_ complex *in situ* in the plasma membrane. We used this information to refine the structural model of the 2:2 complex and the dimer interface with MD simulations, resulting in a comprehensive picture of the early events of MET receptor activation by *L. monocytogenes*. Our approach is generally applicable to distinguish between conflicting structural models of ligand-receptor complexes or to scrutinize putative complex structures predicted by low-resolution experiments or computational tools such as AlphaFold ^20^ and RoseTTaFold ^21^.

## Results

### Binding of InlB promotes an extended conformation of the MET ectodomain

The heavily glycosylated MET ECD consists of six domains: the Semaphorin (Sema), the plexin-semaphorin-integrin (PSI), and four repeated immunoglobulin-like IPT1-IPT4 (Ig-like, plexins, transcription factors) domains. The intracellular part consists of the JM and the tyrosine kinase (TK) domain, and is connected to the ectodomain by a single TM helix ^10,22^ (**Figure 1A**).). Both HGF and InlB bind to the Sema domain, and InlB additionally interacts with the IPT1 domain of the receptor^17,23,24^.

**Figure 1:**
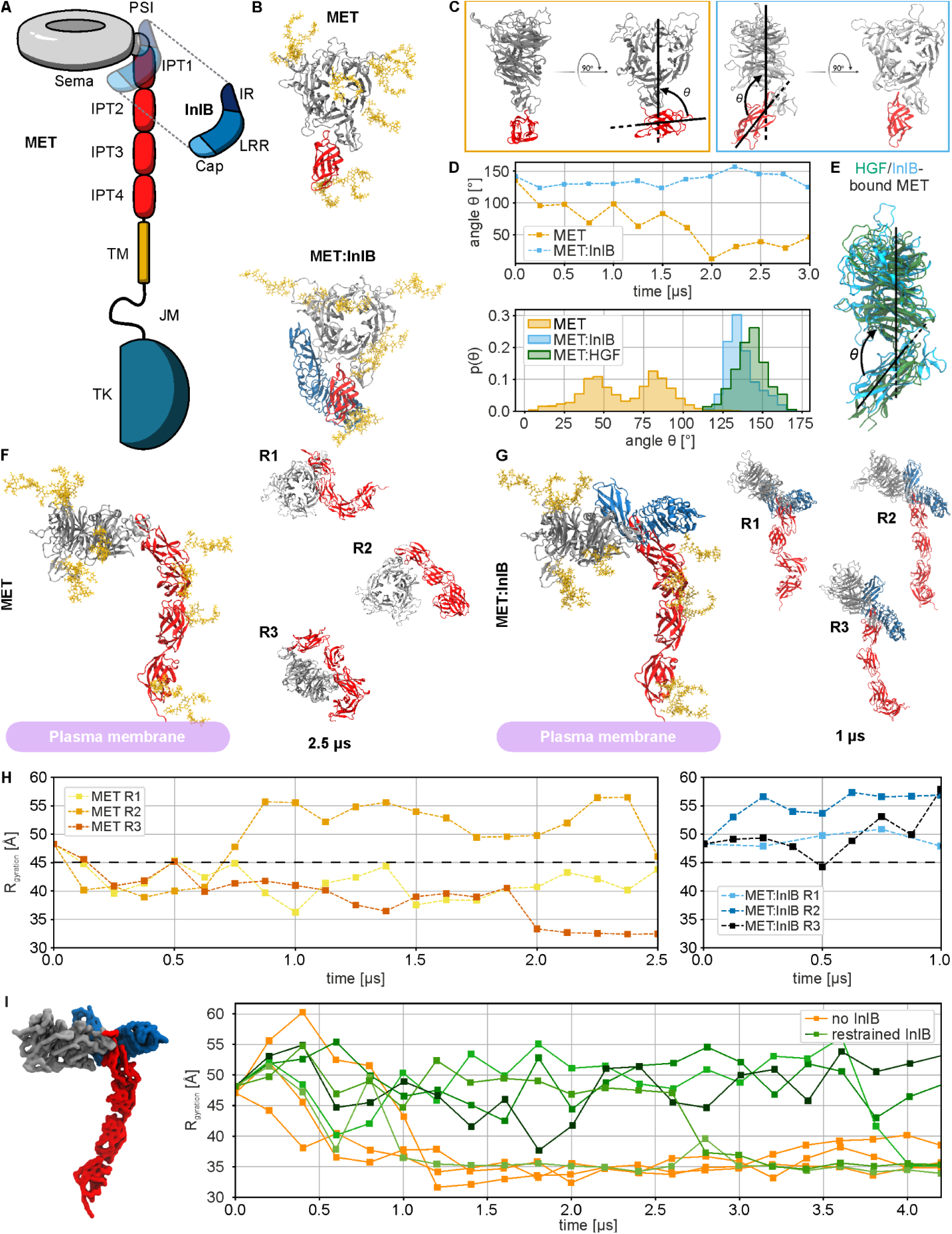
Structural characterization of MET and MET:InlB_321_ obtained with multi-scale MD simulations. (**A**) Schematic representation of the MET receptor bound to InlB_321_. The ligand is represented transparently on the receptor structure. (**B**) Renders of the N-glycosylated MET upper ectodomain system in isolation (top row, MET) and bound to InlB_321_ (bottom row, MET:InlB_321_). InlB is shown as a blue cartoon; Sema and PSI are in silver, the IPT1 domain in red, and the glycans in ochre yellow; water and ions not shown for clarity). (**C**) Side and front views of the closed and open conformations (coloring as in **B,** InlB and glycans not shown for clarity) of MET (orange frame) and MET:InlB_321_ (blue frame). The axes that define the angle θ are reported on the side view renders. (**D**) Time series of the θ angle of MET and MET:InlB models (top panel) and histograms of the θ angle calculated from simulations of the MET:InlB_321_ model, the MET model, and the monomers in the MET dimer in complex with its endogenous ligand HGF (based on PDB 7MO7) (bottom panel). (**E**) Render of one of the monomers involved in the MET:HGF dimer (based on PDB 7MO7) aligned to the InlB-bound MET upper ectodomain model. (**F**) Left: Render of the entire N-glycosylated MET ectodomain model in isolation (Sema and PSI are in silver, the IPT1-4 domains in red and the glycans in ochre yellow; water and ions not shown for clarity). Right: Ectodomain configurations obtained by three replicas (R1-R3), each simulated for 2.5 µs (Sema and PSI in silver, IPT1-4 domains in red; water, ions, and glycans not shown for clarity). (**G**) Left: Render of the N-glycosylated MET entire ectodomain model bound to InlB_321_ (Sema and PSI are in silver, the IPT1-4 domains in red, the glycans in ochre yellow and the InlB_321_ in blue; water and ions not shown for clarity). Right: Ectodomain configurations obtained by three replicas (R1-3), each simulated for 1 µs (Sema and PSI in silver, IPT1-4 domains in red; water, ions, and glycans not shown for clarity). (**H**) Radius of gyration (R_g_) computed on the Cα atoms of the replicas of the MET entire ectodomain model (yellow to red) and of the MET:InlB_321_ entire ectodomain model (blue to black). The black dashed horizontal line at 45 Å indicates the threshold between extended (R_g_ > 45 Å) and collapsed conformations. (I) Left: render of the MET:InlB_321_ model in quasi-atomistic resolution. Sema and PSI are shown in silver, IPT1-4 in red, and InlB in blue. Right: radius of gyration of the quasi-atomistic models of MET (orange) and MET:InlB_321_ (green shades).

We first explored how the binding of the invasion protein InlB affects the structural dynamics of the MET ectodomain. We modeled the upper ectodomain, comprising the Sema, PSI, and IPT1 domains (**Figure 1A**), and ran atomistic MD simulations. To identify the consequences of InlB binding, we compared the dynamics of ectodomain fragments in isolation and in complex with InlB (**Figure 1B**). We chose a minimal version of InlB, InlB_321_, which comprises a cap, a leucine-rich repeat and an inter-repeat region, and which activates MET signaling ^25^.

The simulations revealed that the binding of InlB causes a dramatic reduction in the flexibility of the upper ectodomain of MET. In the absence of InlB_321_, IPT1 explores different orientations with respect to the Sema domain (**Figure 1C**, **Supplementary Fig. 1A**). In particular, PSI acts as a lever between the Sema and IPT1 domains, mediating the interactions between the two domains (**Supplementary Fig. 1B**). To quantify MET structural dynamics, we introduced the angle θ, defined as the angle formed between the Sema and IPT1 domains (**Figure 1C**). In the isolated upper ectodomain, the angle value quickly decreased, corresponding to a structural closing of the IPT1 domain on the Sema (**Figure 1C,D**). In the complex, instead, InlB prevents the Sema and IPT1 domains from closing onto each other. The angle describing the opening between the two domains converges to an average value of θ = 135° (**Figure 1C,D**).

Surprisingly, the conformation assumed by the upper ectodomain of MET in the complex with InlB is very similar to the one of MET in the complex with the endogenous ligand HGF (**Figure 1D,E**). Despite the remarkably different binding modes of these two ligands, the angle formed by the Sema and IPT1 domains in both structures is approximately θ = 135°. Moreover, the equilibrated model of the isolated MET upper ectodomain aligns with the crystal structure of HGF beta-chain in complex with MET ^24^ (PDB 1SHY, see **Supplementary Fig. 1C**).

The MD simulations revealed that the binding of InlB controls the overall conformation of the MET ectodomain. In the absence of the ligand, in three independent MD simulations the chain of IPT domains slowly deviated from a linear arrangement, forming a very compact conformation of the ectodomain (**Figure 1F,H**). In 2 out of 3 replicas, the Sema domain moved close to the terminal IPT4 domain and, therefore, close to the membrane. In contrast, all three MET:InlB complex replicas maintained a stable extended conformation (**Figure 1G,H**). The receptor maintained an upright conformation perpendicular to the plane of the membrane.

We additionally generated a quasi-atomistic model ^26^ of the full MET ectodomain with and without bound InlB (**Figure 1I, Supplementary Fig. 2A)**. This slightly less detailed model enables us to simulate the dynamics of the MET:InlB complex at a reduced computational cost, allowing us to accumulate more statistical evidence.

When bound, the InlB kept the receptor’s stalk extended for up to 3 μs in 4 out of 5 replicas (**Figure 1I** and **Supplementary Fig. 2B,C**) as opposed to the isolated MET which collapsed within the first μs of simulation in both atomistic and quasi-atomistic models.

Both the atomistic and the quasi-atomistic simulations showed that the structural constraints imposed by the binding of InlB on the upper ectodomain propagate non-locally along the whole chain of IPT domains. In the extended conformation of the complex, InlB is exposed and always located at the same height from the membrane, compatible with the formation of a ligand-mediated 2:2 (MET:InlB)_2_ homodimer.

### In situ FRET reports the relative orientation of InlB in (MET:InlB)_2_

We used smFRET to reveal the orientation of two InlB_321_ molecules within the dimeric (MET:InlB)_2_ complex directly in cells. First, we generated two variants of InlB_321_ carrying a single cysteine residue either at position 64 (K64C mutant), termed “H” (head); or at position 280 (K280C mutant), termed “T” (tail) (**Figure 2A**). Using maleimide chemistry, we prepared fluorophore-labeled InlB_321_ variants (ATTO 647N, Cy3B) *in vitro* and determined their degree of labeling (**Supplementary Table 1**). The activity of the fluorophore-labeled InlB_321_ variants was determined from MET phosphorylation in U-2 OS cells using western blotting (**Supplementary Fig. 3**). The affinity of fluorophore-labeled InlB_321_ was previously determined to be very similar to the unlabeled InlB_321_^27^.

**Figure 2:**
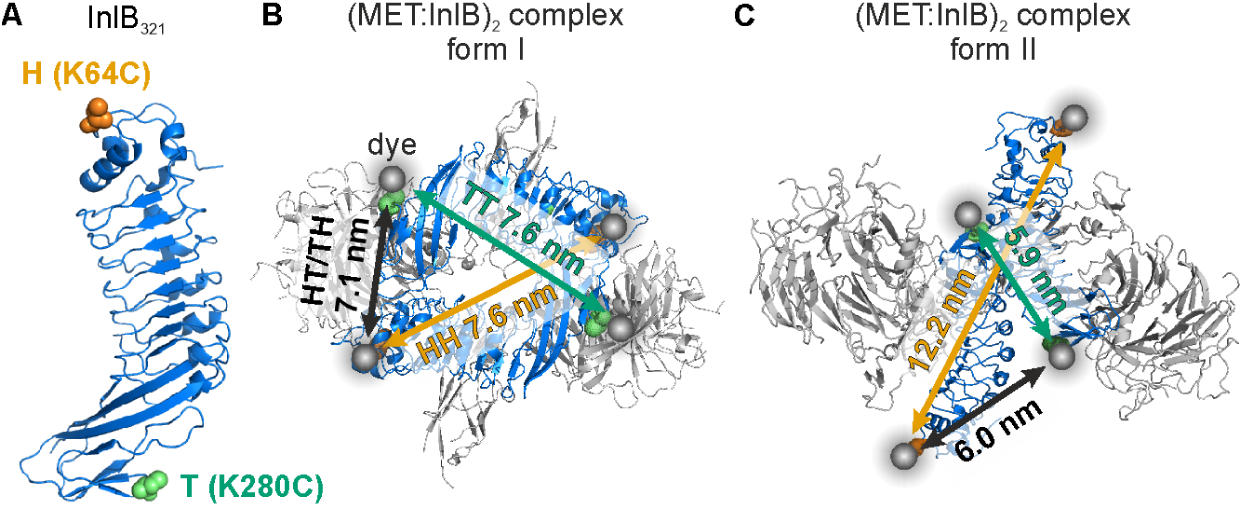
InlB_321_ site-specifically labeled variants in two possible MET:InlB dimer structures differing by the orientation of the MET:InlB monomers (MET in gray and InlB in blue). (**A**) Two InlB variants, K64C (H variant, mutation site highlighted in orange) and K280C (T variant, mutation site highlighted in green), are labeled with donor and acceptor fluorophores for single-molecule FRET. (**B**) Form I assembly of (MET_741_:InlB_321_)_2_ dimer. Donor-acceptor distances between various combinations of two InlB variants were determined by AV simulations, yielding 7.6 nm (T-T), 7.1 nm (H-T/T-H), and 7.6 nm (H-H). (**C**) Form II assembly of (MET_741_:InlB_321_)_2_ dimer. Donor-acceptor distances between two InlB variants were determined by AV simulations, yielding 5.9 nm (T-T), 6.0 nm (H-T/T-H), and 12.2 nm (H-H). The protein structures are adapted from PDB entries 1H6T, 2UZX, and 2UZY, respectively.

Considering the two proposed organizations of (MET:InlB)_2_, two structural assemblies of the dimeric complex (MET:InlB)_2_ are possible: a first one with the form I assembly (PDB 2UZX), and a second one with the form II assembly (PDB 2UZY)^23^. The fluorophore-labeled InlB constructs were designed to distinguish between these two forms by measuring three distances: H-H, H-T/T-H, and T-T (**Figure 2B,C**). The expected donor-acceptor distances for two labeled InlB proteins in the (MET:InlB)_2_ complex were estimated by accessible volume (AV) simulations^28^, yielding distances for the form I of 7.6 nm (H-H), 7.1 nm (H-T/T-H), and 7.6 nm (T-T), and for the form II of 12.2 nm (H-H), 6.0 nm (H-T/T-H), and 5.9 nm (T-T).

Next, we evaluated various adherent cell lines for their suitability to conduct smFRET microscopy of MET receptor complexes. This requires a MET surface density that is sufficiently low for spatial separation of single receptor assemblies with diffraction-limited microscopy. We measured the surface density of MET in various cell lines using *direct* stochastic optical reconstruction microscopy (*d*STORM) ^29^ (**Figure 3A,B** and **Supplementary Table S2**). A first consideration were HeLa cells which are a standard cell line for studies of MET receptor ^30–36^. However, the MET surface expression density in HeLa cells ranged between 6 and 14 clusters/µm^2^ (**Figure 3B**), which is too high for a spatial separation with diffraction-limited microscopy. In single-color imaging experiments, this limitation was bypassed by sub-stoichiometric labeling of MET with InlB_321_ ^37^. However, smFRET experiments require both donor- and acceptor-labeled InlB_321_, and sub-stoichiometric labeling would drastically reduce the probability to detect donor-acceptor labeled (MET:InlB)_2_ dimers (see also **Supplementary Note 1**). From the receptor density quantification (**Figure 3B** and **Supplementary Table S2**), we selected U-2 OS as a cell model for smFRET imaging, because it showed the lowest density of MET on the plasma membrane with 2.8 ± 1.2 clusters/µm².

**Figure 3:**
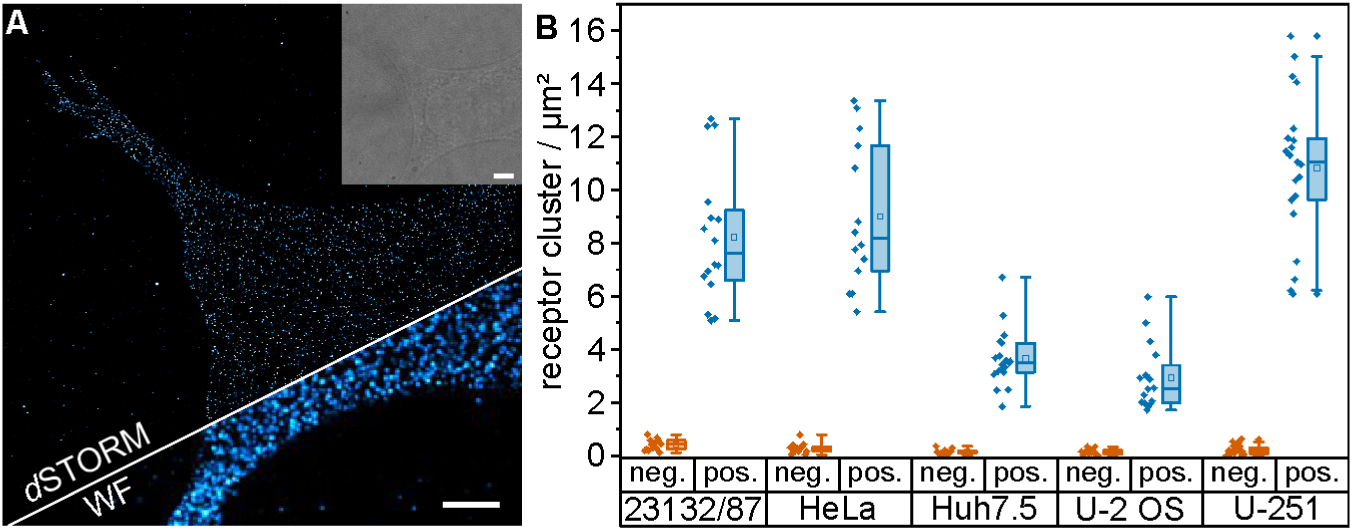
Single-molecule super-resolution imaging of MET receptor densities at the plasma membrane of various cell lines. (**A**) *d*STORM imaging of MET in U-2 OS cells. The super-resolution image (top), the widefield (WF) image (bottom), and the brightfield image (inset) are shown. Scale bars 5 µm. (**B**) MET receptor cluster densities on the plasma membrane of different cell lines. The negative controls (orange) were obtained by incubating the cells with secondary antibodies without prior incubation with primary antibodies. The diamonds represent the receptor densities of single cells. The boxes of the box plots display the 25th and 75th percentile, and the whiskers the 5th and 95th percentiles. In addition, the median (line) and the mean (square) are shown. Receptor densities were obtained from 12-22 cells from at least 3 independent experiments.

Next, we set up an smFRET experiment in fixed cells. We used a widefield microscope operated in total internal reflection (TIR) mode, in order to limit laser excitation close to the glass surface and thus the basal plasma membrane of cells ^38^ (**Supplementary Fig. 4**). In addition, we implemented alternating laser excitation (ALEX), where donor and acceptor fluorophores are excited in an alternating mode using two excitation wavelengths ^38^. ALEX-FRET provides information on both the FRET efficiency (E) and the molecular stoichiometry (S) and enables “molecular sorting” in a 2-dimensional E,S-histogram (**Supplementary Fig. 4C**). After U-2-OS cells were incubated for 15 min at 37 °C with 5 nM of both donor- and acceptor-labeled InlB, they were chemically fixed. Using the fluorophore-labeled H-/T-InlB_321_ variants, samples for the three possible FRET pair combinations H-H, H-T/T-H, and T-T were prepared and measured with ALEX-FRET. FRET was detected for InlB_321_ variant combinations T-T and H-T/T-H. Following accurate correction of experimental FRET data ^39,40^ (see Methods), we generated E,S-histograms and found a single population for both T-T and H-T/T-H (**Figure 4A**). From the E,S-histograms, we extracted FRET efficiencies of 0.863 ± 0.003 (T-T) and 0.560 ± 0.005 (H-T/T-H), respectively (**Figures 4A, Supplementary Fig. 5, Supplementary Note 2, Table 1**). These FRET efficiency values correspond to distances of 4.7 ± 0.4 nm (T-T) and 6.2 ± 0.6 nm (H-T/T-H), respectively. Exemplary FRET time traces for single protein complexes show the expected acceptor photobleaching with a correlated rise in donor intensity (**Figure 4B**). For cells that were labeled with H-/H-InlB_321_, no FRET was detected. However, we detected colocalized fluorescence emission of Cy3B-H-InlB_321_ and ATTO 647N-H-InlB_321_ in single-molecule emission events (**Figure 4C** and **Supplementary Fig. 6**). These observations indicate that the H-H distance in the dimer configuration of MET is outside the distance range of FRET (**Table 1**).

**Figure 4:**
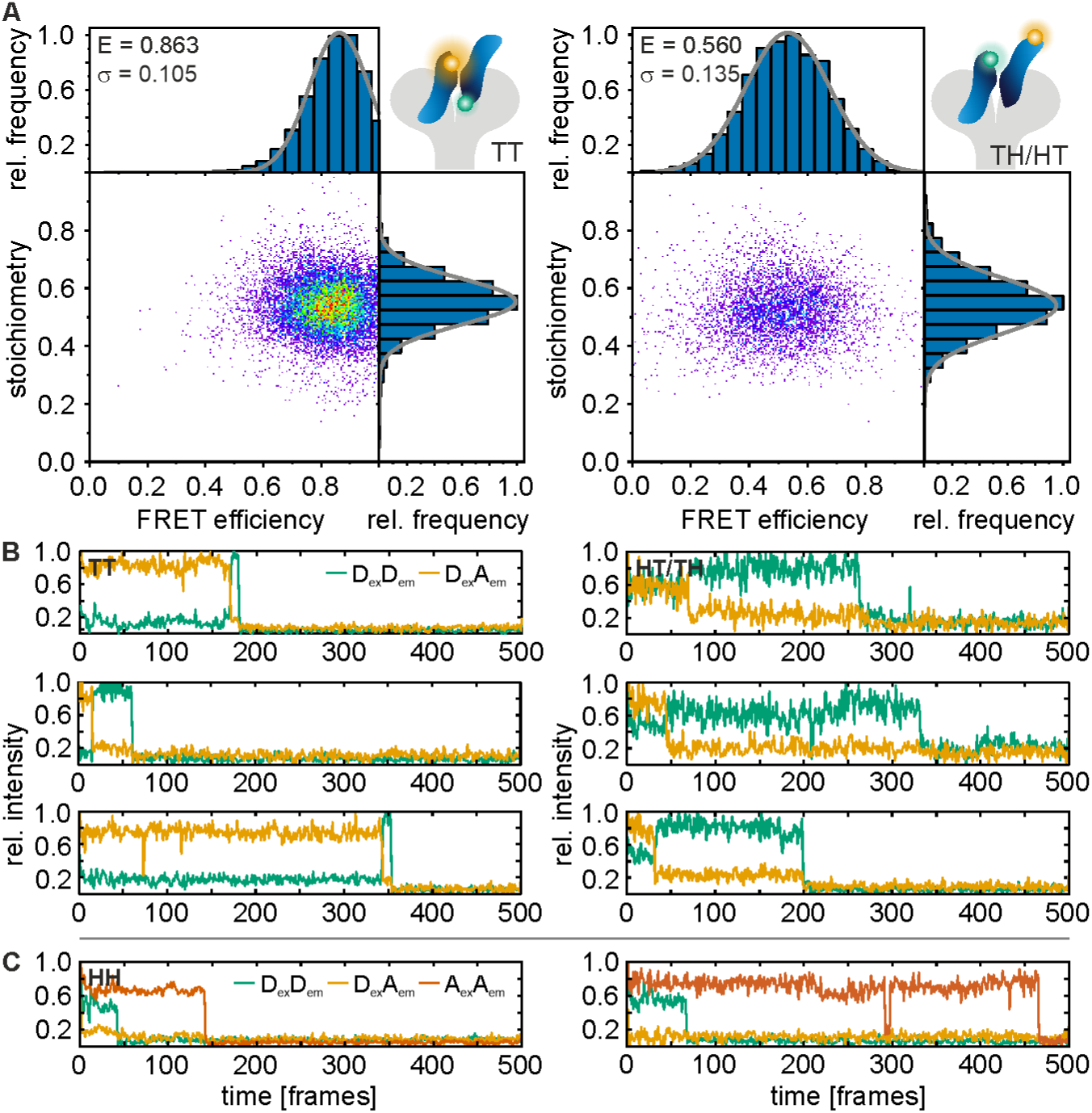
Single-molecule FRET of (MET:InlB_321_)_2_ dimers in U-2 OS cells. (**A**) Left: E, S-histogram for InlB T-Cy3B and T-ATTO 647N (N = 113 smFRET traces from 64 cells); Right: E, S-histogram for InlB T-Cy3B and H-ATTO 647N and InlB H-Cy3B and T-ATTO 647N variants (N = 49 smFRET traces from 39 cells). (**B**) Exemplary smFRET trajectories showing donor (green, D_ex_D_em_) and acceptor (orange, D_ex_A_em_) intensity traces (direct activation of acceptor not shown). **(C)** Exemplary single-molecule intensity traces extracted from colocalized spots showing the fluorescence signal of H-Cy3B and H-ATTO 647N variants showing donor (green, D_ex_D_em_), acceptor (orange, D_ex_A_em_) and direct excitation of the acceptor (red, A_ex_A_em_) fluorescence (see also **Supplementary Fig. 6**). Traces are normalized to 1.

**Table 1:**
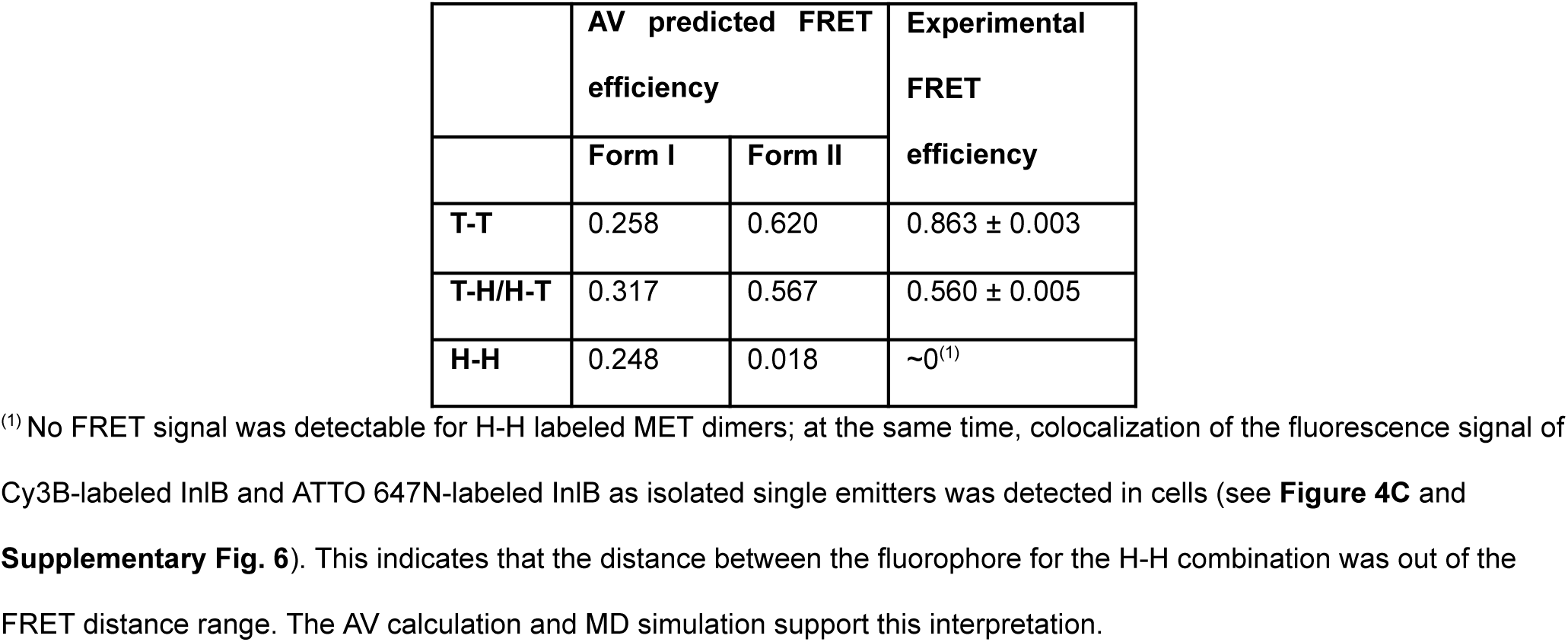
Predicted and experimentally determined FRET efficiencies for different combinations of InlB_321_ variants. Errors are given as standard errors of the mean.

|  | AV predicted FRET efficiency |  | Experimental FRET efficiency |
| --- | --- | --- | --- |
|  | Form I | Form II |  |
| **T-T** | 0.258 | 0.620 | 0.863 ± 0.003 |
| **T-H/H-T** | 0.317 | 0.567 | 0.560 ± 0.005 |
| **H-H** | 0.248 | 0.018 | ~0<sup>(1)</sup> |

<sup>(1)</sup> No FRET signal was detectable for H-H labeled MET dimers; at the same time, colocalization of the fluorescence signal of Cy3B-labeled InIB and ATTO 647N-labeled InIB as isolated single emitters was detected in cells (see **Figure 4C** and **Supplementary Fig. 6**). This indicates that the distance between the fluorophore for the H-H combination was out of the FRET distance range. The AV calculation and MD simulation support this interpretation.

Beyond informing on distances, smFRET can report on structural flexibility. This information can be extracted from the width of the FRET efficiency distribution. We employed photon distribution analysis (PDA), which predicts the theoretical FRET distribution considering the setup-dependent shot-noise and background ^41,42^. PDA yields histograms for the ratio between donor emission and donor-excited acceptor emission that can be compared to experimental data. We found that the PDA histograms are largely identical to experimental data (**Supplementary Fig. 7**), indicating very little structural flexibility in the MET dimer.

A comparison of the smFRET-derived FRET efficiencies to the AV predicted values for the respective fluorophore-labeled InlB variants (**Table 1**) and the finding that the H-H combination did not yield a FRET signal (**Figure 4C**) suggested that the (MET:InlB)_2_ dimer favors a form II assembly in cells. However, the FRET-derived distances are not in a quantitative agreement with the values predicted from the crystal models (**Figure 2B,C**). While the experimental result for the H-T/T-H distance (6.2 nm) is close to the predicted value (6.0 nm), the experimental result for the T-T distance (4.7 nm) is considerably shorter than the predicted value (5.9 nm). This discrepancy is larger than what is expected from the accuracy of AV simulations, and motivated us to investigate the structure of (MET:InlB_321_)_2_ with MD simulations.

### MD simulations of the (MET:InlB)_2_ dimer quantitatively explain the experimental FRET data

We performed atomistic MD simulations of the form II (MET:InlB)_2_ dimer model. We started from the proposed form II structure (PDB 2UZY), containing two copies of the upper ectodomain in complex with InlB (**Figure 5A**). In this model, back-to-back contacts between the two InlB constitute the dimer interface. We then ran three independent replicas each for 600 ns to assess statistical variability. The dimer remained associated in all replicas and sampled only local rearrangements. One replica (R1) remained the closest to the initial starting structures, whereas the other two (R2 and R3) rearranged in a more significant way (**Figure 5B**). Compared to the first replica, which remained close to the initial structural model, the dimeric interface in the third replica was smaller but more compact (**Figure 5C** and **Supplementary Fig. 8**). This interface shows closer contacts between opposite charges and a more compact hydrophobic core.

**Figure 5:**
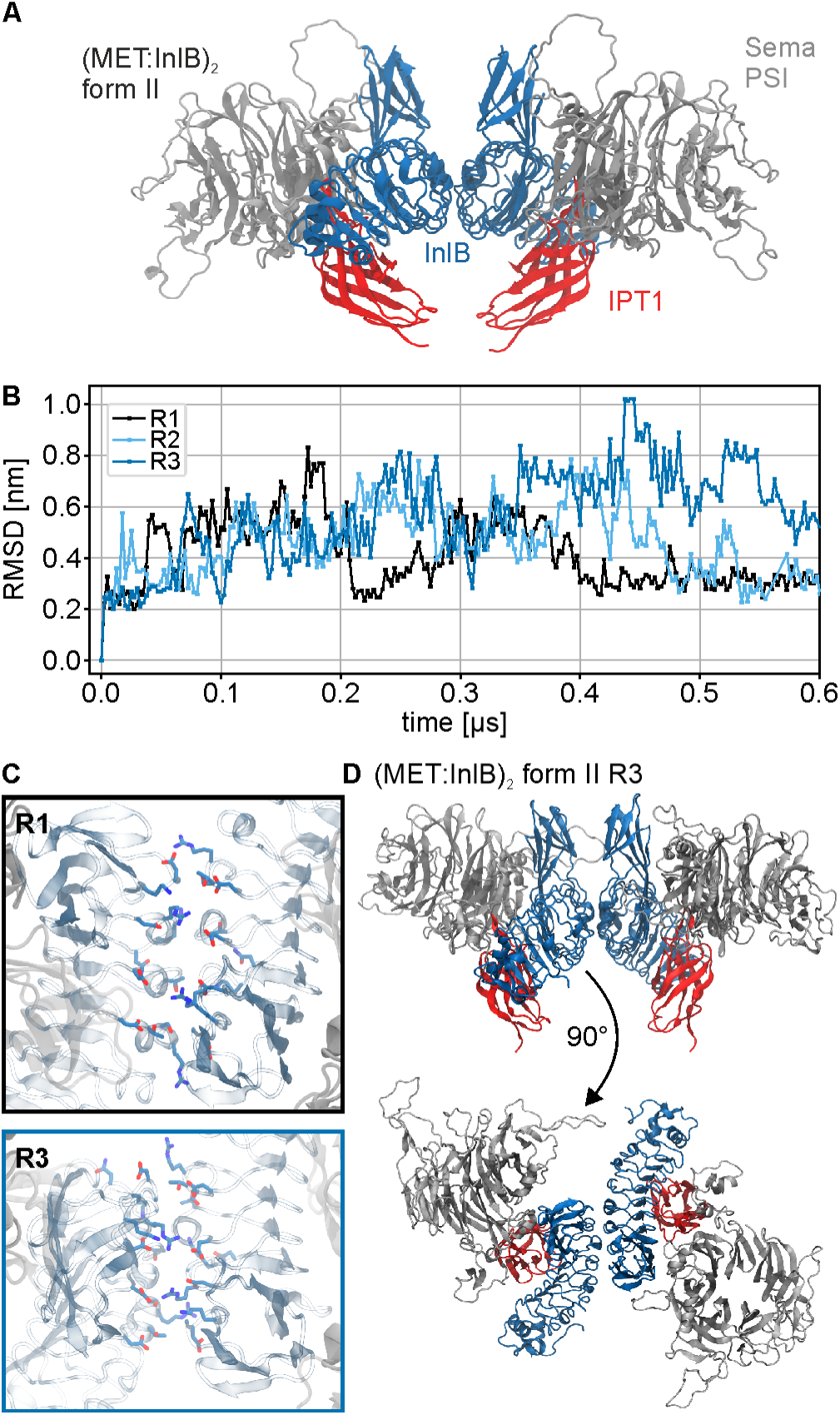
Molecular dynamics simulation of the (MET:InlB_321_)_2_ dimer. (**A**) Renders of the initial form II (MET:InlB_321_)_2_ complex model (Sema and PSI domain of MET in silver cartoon and IPT1 domain in red cartoon; InlB in blue cartoon. Water and ions not shown). (**B**) RMSD time series of the form Il (MET:InlB_321_)_2_ complex model replicas calculated with respect to the first frame. (**C**) Representative assemblies of the two different dimer interfaces explored during the simulations (InlB_321_ in cyan cartoon, MET in silver cartoon, positive side chains in blue, negative side chains in red). Replica 1 explored a broader dimer interface (black frame), while replica 3 explored a more compact one (blue frame; water, ions, and glycans not shown for clarity). (**D**) Render of the proposed antisymmetric dimer structure (explored by R3, compact dimer interface) showing top view (top panel) and side view (bottom panel) (water, ions, and glycans not shown for clarity).

We then calculated distributions of FRET distances for the three replicas (**Table 2**). For this purpose, we used FRETpredict, a novel approach that overcomes limitations in AV calculations. FRETpredict systematically takes into account the protein conformational ensemble and accurately models the conformational ensemble of the fluorophore labels ^43^. The predictions for T-T from the replicas that were locally reorganized (R2 and R3) are incompatible with that from the replica that remained the closest to the initial model. The predicted values for T-T from R2 and R3 went toward a better agreement with the smFRET data. In particular, both T-T and H-T/T-H predicted from R3 are in quantitative agreement with the experimental smFRET data. The results of integrating atomistic MD simulations and smFRET show that *in situ*, the MET:InlB dimer deviates from the crystal form II organization **(Figure 5D**).

**Table 2:** MD-predicted distances for the three FRET dye pairs for each replica (R1, R2, R3) in comparison to the experimental values.

| Dye pair<sup>(1)</sup> | R1 | R2 | R3 | Experiment |
| --- | --- | --- | --- | --- |
| T-T | 3.6 ± 0.1 | 4.1 ± 0.1 | 4.3 ± 0.2 | 4.7 ± 0.4 |
| T-H/H-T | 5.90 ± 0.03 | 5.78 ± 0.04 | 5.75 ± 0.05 | 6.2 ± 0.6 |
| H-H | 10.24 ± 0.02 | 9.81 ± 0.02 | 9.80 ± 0.01 | <sup>(2)</sup> |

<sup>(1)</sup> The errors of the predicted distances were calculated by bootstrapping. This accounts for resampling the FRET efficiency values computed using FRETpredict on the MD trajectories <sup>43</sup>. The bootstrapped efficiencies were converted to FRET distances (using the $R_0$ for the FRET pair), and the standard deviation was computed. For the T-H/H-T distance, we first converted the MD-calculated efficiency into distance prior to bootstrapping. We then applied bootstrapping to the merged distance series and computed the standard deviation.
<sup>(2)</sup> No FRET signal was detected for H-H labeled MET dimers; at the same time, colocalization of the fluorescence signal of Cy3B-labeled InIB and ATTO 647N-labeled InIB as isolated single emitters was detected in cells (see **Figure 4C** and **Supplementary Fig. 6**). This indicates that the distance between the fluorophore for the H-H combination was out of the FRET distance range. The AV calculation and MD simulation support this interpretation.

## Discussion

Despite the key importance of plasma membrane-receptor-mediated cellular events, the structural basis and the dynamics of receptor activation *in situ* is still poorly understood. The analysis of soluble fragments by solution methods to address the oligomeric state and by structural methods like X-ray crystallography or single-particle cryo-electron microscopy (cryo-EM) may lead to incomplete answers. For example, the binding of the GAS6 ligand to the soluble ECD of the receptor tyrosine kinase (RTK) AXL results in a 1:1 complex, although a 2:2 AXL:GAS6 complex is most likely formed on the cell surface ^44^. As a further example, crystallography revealed conflicting structural models for signaling active complexes of fibroblast growth factor (FGF) bound to the ECD of its receptor FGFR, another RTK ^45^. Therefore, reductionists *in vitro* studies often need to be complemented by analysis of receptors in the membrane or a membrane-like environment, like lipid nanodiscs or amphipols that are increasingly used in single-particle cryo-EM. Even then, receptor complexes may require chemical cross-linking to avoid dissociation during purification ^46^.

In this study, we present a comprehensive mechanistic analysis of the early activation steps of the MET receptor upon binding of the bacterial ligand InlB. Two MET:InlB dimer structures (form I, PDB 2UZX; and form II, PDB 2UZY) with contrasting orientation of InlB were proposed ^23^.

As is often the case, deriving the actual quaternary structure from these crystal structures is difficult, because contacts in the crystal may represent either mere crystal packing contacts or physiologically relevant protein-protein interactions. Physiological dimers are usually *C*_2_ symmetric. As both form I and form II of the (MET:InlB_321_)_2_ complex have *C*_2_ point group symmetry, this criterion did not help deciding between both assemblies ^47^. Another criterion used to distinguish crystal-packing contacts from evolved protein-protein interactions is the size of the interface. Form I of the MET:InlB dimer has a substantially larger interface than form II (3700 Å^2^ and 1400 Å^2^, respectively). This is presumably the main reason why the PISA server suggests form I to assemble a stable 2:2 complex in solution, whereas it predicts form II to exist only as 1:1 complex. Experimentally, we never observed dimerization of the MET ectodomain by InlB_321_ in solution ^48^. Therefore, we initially suggested that InlB clusters MET into larger complexes in the plasma membrane without the formation of discrete 2:2 complexes ^23^. Later, we hypothesized that form II could represent a biologically relevant 2:2 complex, although it neither is predicted nor observed to be a stable 2:2 complex in solution ^49^.

The currently available crystal and cryo-EM structures resolve MET only down to IPT2 due to considerable flexibility of the MET stalk region. Hence, the structural organization of the presumed (MET:InlB)_2_ complex in the plasma membrane of cells including the entire MET stalk region remained unclear.

To access the structural organization of membrane receptors *in situ*, we established an integrative structural biology workflow by complementing structural insights with single-molecule experiments, modeling and MD simulations. Based on these findings, we propose a mechanistic model for the early activation of the MET receptor by the bacterial ligand InlB (**Figure 6**).

**Figure 6:**
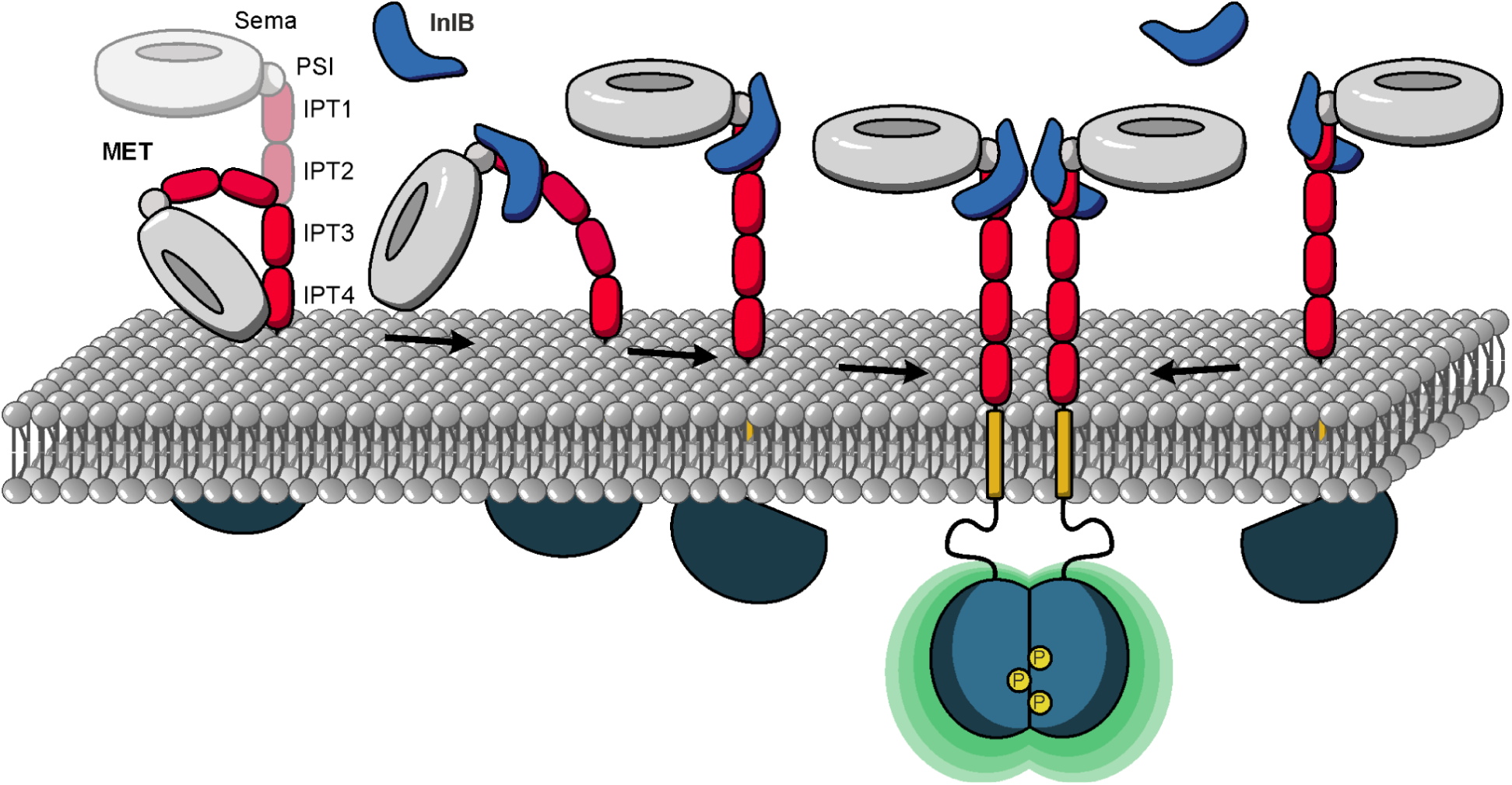
Mechanistic model of MET receptor activation upon InlB binding. In the ligand-free state, the ectodomain of MET shows pronounced flexibility, while the binding of InlB stabilizes an extended conformation. The extended conformation facilitates the association of two MET:InlB complexes to form the signaling-active (MET:InlB)_2_ complex.

Extensive equilibrium MD simulations show that the ectodomain of MET is in a conformational equilibrium between a compact and an extended structure. In the compact conformation, the ectodomain bends significantly, bringing the Sema domain into direct contact with the membrane headgroup region. The ectodomain of integrins, the α-subunit of which is structurally similar to MET, adopts in their inactive form a bent conformation on the membrane surface ^50^ (PDB 3K71), which closely resembles the compact conformation explored by MET’s ectodomain in our simulations (see R2 in **Figure 1F**).

The binding of InlB favors the extended upright conformation of MET’s ectodomain (see **Figure 1G**), which we hypothesize is the signaling-competent monomer. The extended conformation enables back-to-back interactions between two internalin that facilitate the formation of MET dimers. When bound to MET, InlB bridges the Sema domain and the stalk of MET, forcing the structure into a stiff conformation characterized by an openness angle of about θ = 135°. Notably, this is the same angle formed in MET:HGF monomers ^17^, even though the HGF-mediated dimer organization differs significantly from the internalin mediated one. The 135°-conformation appears to be mechanistically critical for the receptor activation.

Informed by crystal structures ^49,51^, we designed a single-molecule FRET experiment to determine the *in situ* structure of the (MET:InlB)_2_ complex (later referred to as *in situ* dimer). Our *in situ* smFRET measurements now unequivocally show that on cells discrete 2:2 MET:InlB complexes do form and they also inform about possible structures of these 2:2 complexes in the native environment of the cell membrane. The smFRET data clearly ruled out the form I assembly in cells under physiological conditions. At the same time, distance information retrieved from smFRET experiments were not in quantitative agreement with those inferred from the crystal structure of form II.

Reconciling this discrepancy required sampling the structural dynamics of the complex with MD simulations and using an accurate model of the smFRET experiment ^43^. Three independent MD replicas showed local rearrangements of the dimer (**Figure 5B**). The third replica (R3) of the form II crystal structure, which aligns well with the T-T and H-T/T-H distances from smFRET experiments, explains how the native (MET:InlB)_2_ dimer attains stability: the dimeric interface was smaller but more compact, with closer contacts between opposite charges and a more compact hydrophobic core (**Figure 5C** and **Supplementary Fig. 8**). The back-to-back arrangement of the InlBs in the identified dimer structure is in accordance with cell-based receptor activation assays using InlB variants intended to block assembly of either form I or form II ^47^ and the increased activity achieved when cross-linking InlB proteins in a similar configuration ^49^. Recurrence of form II in a second crystal form ^52^ further supports our *in situ* (MET:InlB)_2_ dimer model. Lastly, the analysis of the MD trajectories of InlB-bound to the MET ectodomain corroborates the reported lower affinity of the IR-Sema interface compared to the LRR-IPT1 interface ^23^. In particular, we observed that in 1 out of 3 replicas this interface dissociates providing flexibility to the MET stalk. The combination of smFRET experiments and MD simulations elucidated the assembly of the native dimer *in situ*.

A critical step of our analysis was the accurate determination of distances from smFRET data. Two benchmark studies conducted by the smFRET community demonstrated the attainable accuracy of distances from smFRET experiments in DNA and protein samples ^53,54^. Applying these analyses allowed retrieving accurate distances, achieving a precision in quantifying inter-dye distances no greater than 0.2 nm and maintaining an accuracy level below 0.6 nm. The distances obtained from smFRET analysis were confirmed in MD simulations, and allowed the refinement of the structural model of the (MET:InlB)_2_ dimer. In addition, smFRET analysis reports on the structural flexibility of a protein assembly. Photon distribution analysis (PDA) ^41,42^ (**Supplementary Fig. 7**) indicated that the (MET:InlB)_2_ dimer predominantly adopts a single conformation. The H-T/T-H FRET dataset exhibits a slightly larger differential between experimental and simulated data, stemming from the two potential binding positions for these FRET combinations, while the simulated datasets contain only a single-state population (**Supplementary Note 2**).

A recent single-particle cryo-EM study that required forced MET dimerization by addition of a leucine-zipper motif reported two distinct structures for MET dimers bound to the ligands HGF and NK1 ^17^. Interestingly, one HGF ligand was found to be sufficient to dimerize two MET receptors, by binding to two distinct binding sites of MET. This asymmetric 2:1 MET_2_:HGF complex can bind another HGF and assembles into a 2:2 (MET:HGF)_2_ complex. In contrast, binding of the NK1 isoform to MET leads to the formation of a symmetric 2:2 (MET:NK1)_2_ complex, in which the NK1 proteins directly interact in a head-to-tail fashion and form themselves a dimer and that is structurally similar to the (MET:InlB)_2_ dimer ^49,51^. This suggests that (MET:HGF)_2_ and (MET:InlB)_2_ dimers are structurally organized in significantly different ways. However, it is unclear whether different structural arrangements at the level of the ectodomain propagate to the intracellular domain, potentially mediating alternative downstream events.

While our structural model illustrates key events of MET activation, it also sheds light on new exciting questions. The MET ectodomain is glycosylated, but the role played by glycans in its structural dynamics is not understood. In the inactive monomer, the Sema domain is in direct contact with the membrane. Specific interactions between the ectodomain and lipid headgroups could further modulate the conformational equilibrium between the inactive and active monomers.

In summary, our model provides insights into the structural dynamics of monomeric MET and the dynamic interplay between MET and InlB, and provides a useful methodological framework to study receptor activation and dimerization on the plasma membrane. Our study illustrates once more that crystal structures provide excellent working hypotheses but do not always exactly correspond to the conformation of biomolecular complexes in the cell. Integration of *in situ* single-molecule experiments with dynamical molecular simulations represents a powerful approach to determining the organization of complexes in the cell.

## Materials and methods

### Passivation and functionalization of 8-well chambers for single-molecule experiments

8-well chambers (SARSTEDT AG & Co. KG, Nümbrecht, Germany) were prepared by plasma cleaning with nitrogen for 10 min at 80% power and 0.3 mbar using a Zepto B (Diener Electronic GmbH, Ebhausen, Germany). Next, 2 µL of 0.8 mg/mL RGD-grafted poly-L-lysine-graft-(polyethylene glycol) (PLL-PEG-RGD) (prepared according to ^32^) were diluted in 23 µL of ddH_2_O per well. The chambers were incubated with the PLL-PEG-RGD solution at 37°C for 1 h before drying in a sterile bench at room temperature for 2 h. Cells were seeded on the same day that the PLL-PEG-RGD coating was prepared.

### Cell culture

The human osteosarcoma cell line U-2 OS (CLS Cell Lines Service GmbH, Eppelheim, Germany) was cultivated in high glucose DMEM/nutrient mixture F-12 (DMEM/F12) (Gibco, Life Technologies, Thermo Fisher Scientific, Waltham, MA, USA) with 1% GlutaMAX (Gibco), penicillin (1 unit/mL), streptomycin (1 µg/mL; Gibco, Life Technologies) and 10% fetal bovine serum (FBS) (Corning Inc., Corning, NY, USA) at 37°C and 5% CO_2_ in an automatic CO_2_ incubator (Model C 150, Binder GmbH, Tuttlingen, Germany). The cervix carcinoma cell line HeLa (DSMZ, Braunschweig, Germany), the hepatocellular carcinoma cell line Huh 7.5 (DKFZ, Heidelberg, Germany), and the astrocytoma cell line U-251 (CLS Cell Lines Service GmbH) were cultivated in high glucose DMEM with 1% GlutaMAX (Gibco) and 10% FBS (Corning Inc.) and the gastric adenocarcinoma cell line 23132/87 (DSMZ, Braunschweig, Germany) was cultivated in RPMI medium with 1% GlutaMAX (Gibco) and 10% FBS (Corning Inc.) as described above. Cells were split every 3-4 days.

For *d*STORM experiments, 23132/87, HeLa, Huh 7.5, U-2 OS, and U-251 cells were seeded onto PLL-PEG-RGD-coated 8-well chambers in the respective medium with penicillin (1 unit/mL) and streptomycin (1 µg/mL; Gibco, Life Technologies) at densities between 0.5 x 10^4^ to 2.5 x 10^4^ cells/well. For smFRET measurements, U-2 OS cells were seeded onto PLL-PEG-RGD-coated 8-well chambers (300 µL cell suspension with 1 × 10^4^ cells/well) and grown with penicillin (1 unit/mL) and streptomycin (1 µg/mL; Gibco, Life Technologies) for 3 days. For western blots, 2 × 10^6^ cells were seeded in 10 cm dishes and incubated at 37°C and 5% CO_2_ for 3 days.

### dSTORM experiments

#### Immunofluorescence of MET

Two days after seeding, the medium of the cells was exchanged against serum-free medium and the cells were grown for one further day. For immunofluorescence, cells were washed once with 1x PBS pre-warmed to 37°C. Cells were fixed with prewarmed 4% methanol-free formaldehyde in 1x PBS for 10 min. After washing thrice with 1x PBS, samples were blocked with a blocking buffer (BB) containing 5% (w/v) bovine serum albumin (BSA) (Sigma-Aldrich) in 1x PBS for 1 h at room temperature with gentle shaking. The primary antibody (goat@MET, #AF276, R&D Systems, USA) was diluted in BB to a final concentration of 2 µg/mL and incubated for 2 h at room temperature with gentle shaking. After the incubation, the cells were washed three times with 1x PBS. The Alexa Fluor 647 rabbit@goat secondary antibody (2 µg/mL in BB, #A-21446, Invitrogen, Thermo Scientific, Germany) was added to the cells and incubated for 1 h at room temperature with gentle shaking. For negative controls, cells were incubated with secondary antibody only, without primary antibody. After washing three times with 1x PBS, the cells were fixed for 10 min with 4% methanol-free formaldehyde in 1x PBS. Gold beads with a diameter of 100 nm (Nanopartz, USA) were used as fiducial markers.

The gold beads stock solution was vortexed shortly and then sonicated for 10 min. A 1:5 dilution was prepared with 1x PBS and sonicated again for 10 min. The dilution of the fiducial markers was added to the cells and incubated for 15 min. Finally, cells were washed three times with 1x PBS and stored in 0.05% (w/v) NaN_3_ in 1x PBS at 4°C until further use.

#### dSTORM imaging

*d*STORM imaging was performed in an imaging buffer containing *β*-mercaptoethylamine (MEA) as reducing agent and glucose oxidase/catalase as oxygen scavenging system. The imaging buffer containing 10% (w/v) glucose, 100 mM MEA, 50 U/mL glucose oxidase (#G2133-50KU, Sigma), and 5000 U/mL catalase (#C3155, Sigma-Aldrich, Germany) in 1x PBS was prepared freshly before the measurements. The pH was adjusted to 8 with 1 M NaOH.

*d*STORM measurements were performed with an N-STORM microscope (Nikon, Japan). A 647 nm laser was used for the excitation of Alexa Fluor 647 and a 405 nm laser for reactivation. The laser intensity of the 647 nm was set to 0.4 kW/cm². The 405 nm laser was adjusted as necessary to obtain a regular blinking (0-22 mW/cm²). The camera settings were as follows: exposure time 50 ms, EM gain 200, preamp gain 3, frame transfer on, and film lengths 30,000 frames. For each cell line at least three independent experiments were performed.

#### Data analysis

*d*STORM movies were analyzed with the Picasso software ^55^. The point-spread functions of single molecules were localized with Picasso Localize using the following parameters: box side length: 7, min net gradient: 60,000, EM gain 200, baseline 216, sensitivity 4.78, quantum efficiency 0.95, pixel size 157 nm, maximum-likelihood estimation. Drift correction was performed in Picasso Render either with RCC or by picks using the gold beads as fiducial markers. Next, localizations were filtered in Picasso Filter for their standard deviations in x and y direction (0.6-1.6 px). The experimental localization precision was determined in Picasso using the nearest neighbor analysis (NeNA) ^56^. Localizations of the same binding event were linked using six times the NeNA value (or a maximum value of 0.45 px) and 5 dark frames. The number of receptor clusters was determined using the density-based spatial clustering and application with noise (DBSCAN) algorithm ^57^. A radius of two times the NeNA value (or a maximum value of 0.15 px) and a minimum number of 10 localizations were set. The cluster number divided by the cell area (determined in Fiji) yielded the MET receptor cluster density.

### Single-molecule FRET with alternating laser excitation

#### Generation of site-specifically labeled InlB variants

InlB_321_ (comprising amino acids 36-321 of the full-length InlB) was produced by fusing it with a cleavable glutathione-S-transferase (GST) protein using the tobacco etch virus (TEV) protease ^37^. To prevent the formation of unwanted disulfide bonds, a C242A mutation was introduced. This mutation does not affect the binding of MET ^25^. Two InlB variants were generated and the respective mutation K64C (H) or K280C (T) as well as the C242A mutation were introduced into the pETM30 vector using the QuikChange® mutagenesis kit (Stratagene) ^23^. *Escherichia coli* BL21-CodonPlus(DE3)-RIL cells transformed with the vector were cultured in lysogeny broth (LB) medium supplemented with kanamycin and chloramphenicol at 37°C until reaching an optical density at 600 nm of 0.6. Following induction with 0.1 mM isopropyl βD-1-thiogalactopyranoside, InlB_321_ variants were expressed overnight with shaking at 20°C. The cells were harvested through centrifugation and lysed. After centrifugation, the lysate was applied to a glutathione sepharose affinity matrix equilibrated in 1x phosphate buffered saline (PBS). The resin was washed with 1x PBS and TEV protease cleavage buffer, and then resuspended in TEV cleavage buffer. TEV protease and dithiothreitol (DTT) were added and incubated at room temperature overnight for cleaving InlB_321_ from the GST tag. InlB_321_ was purified further using anion exchange chromatography. Specifically, InlB_321_ was loaded onto a Source Q 15 column equilibrated with 20 mM Tris buffer pH 7.5 and eluted with a linear gradient of salt concentration (up to 300 mM NaCl).

#### Sample preparation

Three days after seeding, U-2 OS cells were rinsed with 400 µL prewarmed, serum-free DMEM/F12 and then starved for 2 h in serum-free DMEM/F12 at 37°C and 5% CO_2_. For ligand stimulation, Cy3B- and ATTO 647N-labeled InlB_321_ variants (InlB_321_-H or InlB_321_-T) were added to a final concentration of 5 nM per InlB variant. As controls, only one InlB_321_ variant was used. Cells were incubated with the ligand for 15 min at 37°C. Immediately after stimulation, cells were washed once using 200 µL/well of prewarmed 0.4 M sucrose solution in 1x PBS (diluted from 10x stock, #14200067, Gibco), followed by fixation for 15 min at room temperature using a solution consisting of 4% formaldehyde (Thermo Scientific) and 0.01% glutaraldehyde (Sigma-Aldrich) in 0.4 M sucrose and 1x PBS. Subsequently, cells were rinsed three times using 300 µL 1x PBS.

To reduce photobleaching during single-molecule measurements, an oxygen scavenging buffer (300 µL/well) was employed which was prepared freshly before each measurement: glucose oxidase from *Aspergillus niger* type VII (0.009 U/µL; Sigma-Aldrich), catalase from bovine liver (594 U/mL; Sigma-Aldrich), glucose (0.083 M; Sigma-Aldrich), and Trolox (1 mM; Sigma-Aldrich) ^58,59^.

#### Setup and data acquisition

Single-molecule FRET measurements were performed on a home-built total internal reflection fluorescence (TIRF) microscope based on an Olympus IX-71 inverted microscope (Olympus Deutschland GmbH, Hamburg, Germany). The excitation light was provided by two lasers (637 nm, 140 mW OBIS and 561 nm, 200 mW Sapphire, both Coherent Inc., Santa Clara, CA, USA). Both laser beams were colinearly superimposed using a dichroic mirror (H 568 LPXR superflat, AHF Analysentechnik AG, Tübingen, Germany). An acousto-optical tunable filter (AOTF; AOTFnC-400.650-TN, AA Opto-Electronic, Orsay, France) selected the excitation light, which alternated between 561 nm and 637 nm. The required timing was achieved by means of two digital counter/timer and analog output devices (NI PCI-6602 and NI PCI-6713, National Instruments, Austin, TX, USA). To spatially overlay both lasers and clean the beam profiles, the lasers were coupled by a fiber collimator (PAF-X-7-A, Thorlabs, Dachau, Germany) into a single-mode optical fiber (P5-460AR-2, Thorlabs) and subsequently re-collimated to a diameter of 2 mm (60FC-0-RGBV11-47, Schäfter & Kirchhoff, Hamburg, Germany). The collinear beams were then directed to a 2-axis galvo scanner mirror system (GVS012/M, Thorlabs) where electronic steering, controlled by an in-house Python script, allowed switching between wide-field illumination, steady-state and circular TIRF, and HILO (highly inclined and laminated optical sheet) modes of operation. The excitation beams were then directed through two telescope lenses (AC255-050-A-ML and AC508-100-A-ML, Thorlabs) which focused the beams onto the back focal plane of the objective (UPlanXApo, 100x, NA 1.45, Olympus Deutschland GmbH). In a filter cube, which directs the beam into the objective, two clean-up and rejection bandpass filters together with a dichroic mirror were installed (Dual Line Clean-up ZET561/640x, Dual Line rejection band ZET 561/640, Dual Line beam splitter zt561/640rpc, AHF Analysentechnik AG). A nosepiece stage (IX2-NPS, Olympus Deutschland GmbH) provided z-plane adjustment and minimized drift during the measurements.

Fluorescence emission was collected through the same objective and passed the dichroic mirror towards the detection path. An Optosplit II (Cairn Research Ltd, UK) was used to split the fluorescence light around 643 nm into two channels using a beam splitter together with two bandpass filters (H643 LPXR, 590/20 BrightLine HC, 679/41 BrightLine HC, AHF Analysentechnik AG). The two spatially separated donor and acceptor channels were simultaneously detected on an EMCCD camera (iXon Ultra X-10971, Andor Technology Ltd, Belfast, UK). The setup achieved a total magnification of 100x, resulting in a pixel size of 159 nm. The µManager software ^60^ captured 1,000 frames with the following settings: exposure time 100 ms, EM gain 150, preamp gain 3x, readout rate 17 MHz, image size 512 x 256 pixel, and activated frame transfer. Bright field images of the cells were taken after each measurement. The excitation laser wavelengths were alternated between 561 nm and 637 nm for a duration of 100 ms each. For each sample, four independent experiments were performed. To align both channels, daily measurements of 100 nm TetraSpeck^TM^ microspheres (Invitrogen, Thermo Fisher Scientific, Waltham, MA, USA) on coverglass were conducted for 100 frames without alternating lasers.

#### Data analysis

Single-molecule FRET movies were analyzed using the iSMS software ^39^. The 561 nm and 637 nm excitation channels were aligned with the default settings of the autoalign ROIs tool. FRET pairs were detected averaging the intensity of all 1,000 frames. Initially, we considered every donor and acceptor position as a potential FRET pair. We manually selected FRET traces based on two criteria: an increase in donor intensity upon photobleaching of the acceptor and single-step photobleaching in both the donor and acceptor channels to ensure that only a single donor-acceptor fluorophore pair was present.

Selected smFRET intensity traces were corrected in iSMS for donor emission leakage into the acceptor channel (α), acceptor direct excitation by the donor excitation laser (δ), and different detection efficiencies and quantum yields of donor and acceptor (γ) ^39^. The iSMS software determined α, δ, and γ trace-wise. The mean correction factors were applied to the data within iSMS. In addition, we manually calculated the β-correction factor which normalizes for different excitation intensities and cross-sections of donor and acceptor.

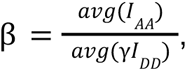

where *I_AA_* represents the emission intensity of directly excited acceptor and *I_DD_* denotes the donor emission intensity from direct excitation. The FRET efficiencies and stoichiometries were determined following a published protocol ^53^ and computed with OriginPro (OriginLab Corporation, Northampton, MA, USA):

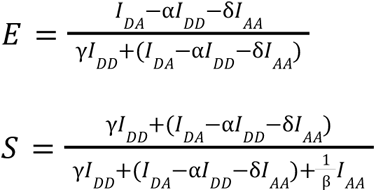

Here, *I_DA_* is the acceptor intensity when the donor is excited. The calculated FRET efficiencies were histogrammed and the distribution for each condition was fitted with a Gaussian distribution to obtain the FRET efficiency for the respective condition. The distances *R* between donor and acceptor fluorophores were calculated from these FRET efficiencies.

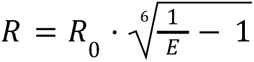

Here, *R*_0_ is the fluorophore-pair-specific Förster radius.

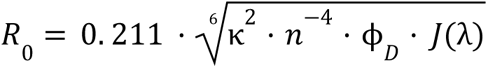

For the orientation factor κ^2^, free rotation of the fluorophores was assumed, therefore κ^2^ – 2/3. The refractive index *n* of the imaging solution was measured to be 1.34. The quantum yield ϕ*_D_* of the donor is given by the fluorescence decay rate *k_F_* and the fluorescence lifetime τ_*L*_.

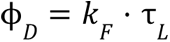

For Cy3B, ϕ_*D*_ was calculated from the fluorescence decay rate (0.239 ns^-1^; calculated using quantum efficiency and lifetime from Cooper et al. ^61^) and the lifetime of the donor determined by time-correlated single photon counting (TCSPC) for each Cy3B-labeled InlB_321_ variant (T variant: *τ_L_* = 2.6 ns, H variant: *τ_L_* = 2.5 ns). The fluorescence lifetimes were measured using a PicoHarp 300 system (Picoquant, Berlin, Germany) in combination with a pulsed 485 nm laser for excitation. Finally, *J*(λ) represents the overlap integral of Cy3B emission and ATTO 647N absorption (5.8·10^15^ M^-1^ cm^-1^ nm^4^; FPbase ^62^). The Förster radius *R*_0_ calculated for the T variant is 6.34 nm and for the H variant 6.30 nm. The standard deviation of the distance *R* was obtained by fitting a Gaussian distribution to the histogram of all individual distances.

The photon distribution analysis (PDA) was conducted using software provided by the group of Claus Seidel (https://www.mpc.hhu.de/software/pda). The underlying model for this analysis is derived from Antonik et al. ^41,42^. Emissions from both, donor and acceptor, under donor excitation, were converted into a 2D histogram. Subsequently, this histogram was imported into the Tatiana software for further analysis. The fitting of the ratio between donor and acceptor emission followed Antonik et al.’s approach, employing free fit parameters, except for two fixed parameters: the number of limited width states and dynamic states, both set at 1.

#### Donor-acceptor distance estimation by AV simulation

To estimate the distances between donor and acceptor in the (MET:InlB_321_)_2_ complex for different InlB variants, we applied accessible volume (AV) simulations ^63^. AV simulations predict the allowed average distances between donor and acceptor dyes. It was achieved by the FRET Positioning and Screening (FPS) software ^28^ using the parameters summarized in **Table 3**. The FRET-averaged distances are shown in **Figure 2**.

**Table 3:** AV simulation parameters used for donor-acceptor distance estimation within the (MET:InlB_321_)_2_ complex. The linker is simplified as a cylinder model; the length and width represent the height and radius of the cylinder. The dye is simulated as an ellipsoid using the 3AV model. The radii R_1_, R_2_, and R_3_ describe the dye ellipsoid. Linker and dye dimensions were taken from Klose et al. ^64^ for Cy3B maleimide and from Claus Seidel (Heinrich Heine University Düsseldorf) for ATTO 647N maleimide.

| | Linker length [Å] | Linker width [Å] | $R_1$ [Å] | $R_2$ [Å] | $R_3$ [Å] |
| --- | --- | --- | --- | --- | --- |
| ATTO 647N maleimide | 21.0 | 4.5 | 7.15 | 4.5 | 1.5 |
| Cy3B maleimide | 18.5 | 4.5 | 3.4 | 8.2 | 3.0 |

### Western blots

U-2 OS cells were rinsed with 5 mL serum free DMEM-F12 per dish and then starved in 10 mL serum-free DMEM-F12 medium for at least 8 h at 37°C. After starvation, the cells were stimulated at 37°C for 15 min with 2 mL of 5 nM InlB_321_ variant (Table S1) or 1 nM HGF (# 100-39H, PeproTech, Hamburg, Germany) in serum-free DMEM-F12. For the resting condition, cells were only treated with serum-free DMEM-F12 medium for 15 min. Then cells were rinsed with 10 mL ice-cold 1x PBS and kept for 2 min on ice. PBS was then removed and 80 µL of lysis buffer (Triton X-100 1%, Tris-HCl (pH 7.4) 50 mM, NaF 1 mM, NaCl 150 mM, Na_3_VO_4_ 1 mM, EDTA 1 mM, and ¼ cOmplete Mini, EDTA-free protease inhibitor tablet, Roche for 10 mL) were added per dish and incubated on ice for at least 30 s. The cells were scraped thoroughly to one corner of the dish and collected in ice-cold 1.5 mL tubes.

When all the samples were collected on ice, they were shaken at 750 rpm and 4°C for 5 min and centrifuged at 12,000 rpm and 4°C for 20 min, the supernatants were collected in new tubes and stocked shortly on ice. The concentration of total proteins was determined with the BCA Protein Assay Kit (VWR International GmbH). According to the total protein amount in each sample, 1 M DTT, 5x loading dye (Tris-HCl (pH 6.8) 250 mM, SDS 8% (w/v), bromophenol blue 0.1% (w/v), glycerol 40% (v/v)), and ddH_2_O were mixed so that the protein amount was 30 µg protein and the final concentrations were 100 mM DTT and 1x loading dye. The samples were stored at −20°C until further use. For sodium dodecyl sulfate-polyacrylamide gel (SDS-PAGE), samples were heated to 95°C for 5 min before cooling down on ice. Each pocket of the SDS-PAGE gel (cat. 4561094, Bio-Rad, Hercules, CA, United States) was filled with 35 µL sample or 6 µL PageRuler (cat. 26617, Thermo Fisher Scientific). Gel electrophoresis was performed in running buffer (Tris base 25 mM, glycine 192 mM, SDS 3.46 mM in ddH_2_O) at 170 V for around 45 min. The protein was transferred from the gel to western blot with an iBlot gel transfer system (cat. IB1001, Invitrogen, Thermo Fisher Scientific) for 7 min. Each blot was blocked with 10 mL 5% (w/v) nonfat dry milk (Cell Signaling Technology, Danvers, MA, USA) in TBST buffer (25 mM Tris base, 150 mM NaCl, and 0.05% (v/v) Tween-20, pH 7.6) for 1 hour at room temperature. After blocking, the blots were washed 3 times with TBST buffer and shaken gently with 5 mL primary antibody (rabbit anti-MET, #4560, Cell Signaling Technology, 1:1000 dilution, or rabbit anti-pMET, #3077, Cell Signaling Technology, 1:1000 dilution, and rabbit anti-actin, #ab14130, abcam, Cambridge, UK, 1:10000 dilution) in 5% (w/v) bovine serum albumin in TBST at 4°C overnight. The excess of primary antibodies was removed by washing 3 times with TBST. The blots were incubated with 10 mL secondary antibody (goat anti-rabbit HRP, cat.111-035-003, Jackson ImmunoResearch, West Grove, PA, USA, 1:20000 dilution in 5% (w/v) BSA in TBST) at room temperature for 3 hours. Afterwards, the blots were rinsed 4 times with TBST and one time with TBS ((25 mM Tris base and 150 mM NaCl, pH 7.6). Every wash step was incubated for at least 5 min. The blots were visualized by CHEMI-only chemiluminescence imaging system (VWR). The quantitative analysis was done in the open-source Fiji software (NIH, USA) ^65^. All chemicals for which the manufacturer was not named were purchased from Sigma-Aldrich.

### Atomistic molecular dynamics of MET upper ectodomain

We modeled the atomistic upper ectodomain (UniProt P08581-1 sequence numbering, residues 43-657) of the MET receptor in isolation and in complex with the InlB_321_ fragment of the InlB protein starting from the crystallographic structure PDB 2UZY ^23^. We modeled the missing residues (UniProt P08581-1 sequence numbering, 92-110, 151-155, 206-209, 302-311, 378-383, 398-406, 411- 413, 628-633) with the MODELLER ^66^ implemented on UCSF Chimera ^67^. We added 8 A2 N-glycans (di-sialylated, bi-antennary complex-type N-glycans) in both models on experimentally determined N-glycosylation sites (UniProt P08581-1 sequence numbering, residue 45, 106, 149, 202, 399, 405, 607, 635) ^68^.

We solvated the systems using CHARMM-GUI in combination with GROMACS ^69^ as MD engine. We minimized the systems using the steepest-descent algorithm for 5000 steps and performed a 125-ps-long NVT (particle amount, volume and temperature are kept constant) equilibration using the Nose-Hoover thermostat with a reference temperature of 310 K (*τ*t = 1 ps). We simulated the systems in the NPT (particle amount, pressure and temperature are kept constant) ensemble for 3 µs each using the Charmm36m forcefield ^70,71^ with TIP3 water model, a reference temperature of 310 K (*τ*t = 1 ps, V-rescale thermostat), a reference pressure of 1 bar (*τ*p = 5 ps, Parrinallo-Rahman barostat) and a NaCl concentration of 0.15 M. For both the Van der Waals (Verlet) and the Coulomb forces (Particle Mesh Ewald) we used a r_cut−off_ = 1.2 nm. We used a 2 fs timestep. We used GROMACS 2021.3 ^69^ for the MET:InlB system and GROMACS 2021.4 ^69^ for the isolated MET model.

### Atomistic molecular dynamics of the entire MET ectodomain

We initially produced two models of the entire ectodomain of the MET receptor (UniProt P08581-1 sequence numbering). The first one spans residues 43-930. The second one, spans residues 1-930, and contains an N-terminal loop of 42 amino acids and a disulfide bond between CYS26 and CYS584 on IPT1, which could affect the ectodomain’s structural dynamics. For the first model we started from the equilibrated upper ectodomain model as described above. In absence of an experimental structure, we built on the AlphaFold prediction ^20^ of the IPT2, IPT3, and IPT4 domains. We used the MET receptor structure reported on the AlphaFold database ^72^ corresponding to the UniProt entry P08581. We trimmed the IPT2-IPT3-IPT4 fragment (UniProt P08581-1 sequence numbering, residues 658-930) of the predicted structure and connected it to the upper ectodomain models (MET and MET:InlB) using UCSF Chimera ^67^. For the second model, we started from the cryo-EM structure PDB 7MO7 ^17^, which consists of the SEMA, PSI, IPT1, IPT2 domains. In absence of an experimental structure, we modeled the IPT3-IPT4 by connecting the IPT3-IPT4 fragment from the AlphaFold prediction ^20^ to the cryo-EM model using UCSF Chimera ^67^.

In the crystal structure of the InlB-bound MET the 43-residue N-terminal loop is not resolved, unlike in the cryo-EM structure, which includes it but does not show InlB binding interfaces. To rule out modeling biases in the MET:InlB_321_ complex, we used the experimentally determined crystal structure, expecting that the presence of the disulfide bond would not significantly affect the system due to the observed stability of the upper ectodomain when simulated bound to InlB_321_

Additionally, the comparison of the isolated ectodomain structural dynamics in the two models showed that the N-terminal loop and the CYS26-CYS584 bond are consistent with a very compact conformation and do not significantly affect the extended-compact dynamics of the ectodomain (**Supplementary Figure 10**). To enable an optimal comparison between the isolated and InlB-bound models, we focused our simulations on the crystal-based models, which are also slightly smaller and enable longer simulated trajectories and therefore better statistics. To account for the impact of N-glycosylation we included 11 A2 glycans in both models (UniProt P08581-1 sequence numbering, residue 45, 106, 149, 202, 399, 405, 607, 635, 785, 879, 930) ^68^.

We prepared the systems using CHARMM-GUI solution builder ^73^ in combination with GROMACS ^69^ as MD engine. We minimized the systems using the steepest-descent algorithm for 5000 steps (for the MET in isolation with N-terminal loop we performed two minimization runs) and performed a 125-ps-long NVT equilibration using the Nose-Hoover thermostat with a reference temperature of 310 K (*τ*t = 1 ps). We simulated the systems in the NPT ensemble for 2.5 µs each using Charmm36m forcefield with GROMACS 2021.4 with TIP3 water model, a reference temperature of 310 K (V-rescale thermostat, *τ*t = 1 ps), a reference pressure of 1 bar (Parrinallo-Rahman barostat, *τ*p = 5 ps) and a NaCl concentration of 0.15 M. For both the Van der Waals (Verlet) and the Coulomb forces (Particle Mesh Ewald) we used a r_cut−off_ = 1.2 nm. We chose a 2 fs timestep. Visual inspection of the trajectories revealed that the isolated systems had a similar flexibility, showing collapse of the receptor stalk in 2 out of 3 replicas (see **Supplementary Fig. 9A,B**). To quantify the extension of the ectodomain we exploited the radius of gyration (R_g_). We calculated the R_g_ using the MDAnalysis function radius_of_gyration ^74^.

### Quasi-atomistic molecular dynamics of the entire MET ectodomain

We produced quasi-atomistic coarse-grained (CG) models of MET in isolation and in complex with InlB_321_ using the Charmm-GUI Martini maker webserver selecting MARTINI 3 forcefield ^26^ with elastic network. We removed the interdomain elastic bonds and, to maintain the ligand in place, we applied harmonic restraints between the InlB and the binding interfaces on both SEMA and IPT1 domains using a distance threshold of 2 nm between the backbone beads (**Supplementary Fig. 2A)**. We performed both operations using custom python scripts.. Following the same procedure detailed above, we obtained a coarse-grained model of the MET in isolation. For this model, we removed the interdomain elastic bonds using a custom python notebook and modified the bond type from the standard type 1 to 6.

We then used the *insane.py* script ^75^ to solvate the systems. We used a NaCl concentration of 0.15 M with neutralizing ions. Using GROMACS ^69^, we ran two rounds of minimization using the steepest descent algorithm. The first ran until convergence, the second for 6000 steps. Then, we ran the NPT equilibration at a reference temperature of 310 K (V-rescale thermostat, *τ*t = 1 ps) and a reference pressure of 1 bar (Berendsen barostat, isotropic coupling type, *τ*p = 5 ps). We simulated the systems in the NPT ensemble at a reference temperature of 310 K (V-rescale, *τ*t = 1 ps) and a reference pressure of 1 bar (Parrinello-Rahman barostat, isotropic coupling type, *τ*p = 12 ps). Coulombic interactions were treated with PME and a cut-off of 1.1 nm, as for van der Waals interactions. We used a time step of 20 fs. For the MET:InlB_321_ complex system we used GROMACS 2022.6 ^69^, for the MET in isolation we used GROMACS 2020.5 ^69^.

### Definition of θ angle

We computed θ as defined by two vectors describing the relative orientation of the Sema and the IPT1 domains. The first vector connects the centers of mass of two groups of atoms on the upper and lower side of the Sema domain (182-200 and 464-479); the second vector connects two groups of atoms at the opposite sides of the IPT1 cylinder (561-657 and 655-657). We calculated the value of θ from the simulated atomistic trajectories using custom-written code in the Python packages NumPy ^76^ and MDAnalysis ^74^.

### Molecular dynamics of the (MET:InlB)***_2_*** upper ectodomain dimer

We created an atomistic model of the MET:InlB dimer in the form II as reported by PDB 2UZY ^77^ by aligning two copies of the NVT equilibrated model described in the “***Atomistic molecular dynamics of MET upper ectodomain***” section. We then simulated 3 atomistic replicas of this model. Firstly, we solvated the system with TIP3P water using GROMACS ^69^. We minimized the systems using the steepest-descent algorithm for 5000 steps and performed a 125-ps-long NVT equilibration using the Nose-Hoover thermostat with a reference temperature of 310 K (*τ*t = 1 ps). We simulated each replica in the NPT ensemble for 0.6 µs using the Charmm36m forcefield, a reference temperature of 310 K (*τ*t = 1 ps, V-rescale thermostat), a reference pressure of 1 bar (*τ*p = 5 ps, Parrinallo-Rahman barostat) and a NaCl concentration of 0.15 M. For both the van der Waals (Verlet) and the Coulomb forces (Particle Mesh Ewald) we used a r_cut−off_ = 1.2 nm. We chose a 2 fs timestep. We used GROMACS 2021.4.

### Prediction of FRET distances from atomistic MD simulations

We predicted smFRET distances from the atomistic MD simulations of the MET:InlB form II dimer using the Python package FRETpredict ^43^. This method uses rotamer libraries of FRET dyes superimposed to protein structures or trajectories to predict the FRET efficiency distributions ^43^. It considers the structural dynamics of the FRET dyes and their linkers. We adapted the tutorial jupyter notebooks (download at https://github.com/KULL-Centre/FRETpredict) for our experiments. As the rotamer libraries for our dye pair were not available, we performed ∼ 1.2 µs atomistic MD simulations for each dye in solution. To correctly reproduce the dynamics of the dyes, we used the CHARMM-DYES forcefield, which includes optimized parameters for our FRET dye pair ^78^. We employed CHARMM-DYES combined with the same solvation conditions used in the simulations of the 2 alternative dimer models. The CHARMM-DYES forcefield did not include parameters for the maleimide ring used in experiments nor the thioester bond between the linker and the cysteine residue. Therefore, to approximate the experimental linker length and flexibility, we used a C4 linker where the first two dihedrals were disregarded to account for the stiffness of the missing ring while retaining almost the same bond length. We used the calculated rotamer libraries to perform the FRET efficiency prediction. To account for the position of the linker as attached to the S atom of the cysteine, we set the offset for the rotamer placement on the Cγ atom of the corresponding residue.

The FRET signal produced by the dye pair in T-H/H-T arrangements is indistinguishable in the experiments due to the isotropic character of the dimerization process after InlB-treatment. We, therefore, averaged the predictions of distributions of T-H and H-T. The position of the residues on the InlB enabled us to use the k2 approximation, which allowed us to obtain the efficiency values predicted using the static calculation ^43^. We calculated the distributions of the T-T and T-H/H-T efficiencies for the 3 different replicas. We assessed the local convergence of the replicas by calculating the RMSD of the Cα and Cβ atoms in the InlB-InlB dimer. All FRET predictions were obtained on the last 200 ns of each replica. We estimated the standard deviation of the FRET predictions by applying bootstrapping on the time series of the predicted FRET signal. To perform this task, we used a *pandas* function *Series* in combination with the *sample* function.

## Data availability

Single-molecule imaging data is available on request from the authors. The MD simulation data and parameter files and analysis code are freely available on Zenodo (DOI: 10.5281/zenodo.11202330).

## Author contributions

M.H. and R.C. designed the project. H.H.N. designed the FRET study, and together with D.H. and D.M.F., provided labeled proteins. P.F. assisted with cell culture. H.-D.B. designed the optical setup for smFRET. Y.L. and M.S.D. performed microscopy experiments, and together with M.H. analyzed and interpreted the data. S.A. and G.J.H. performed MD simulations, and together with R.C. analyzed and interpreted the data. M.H., R.C., Y.L., S.A., H.H.N. and M.S.D. discussed the data and wrote the manuscript.

## Materials and correspondence

Correspondence and material requests should be addressed to.

## Acknowledgement

We thank Björn Hellenkamp for helpful suggestions for the AV simulations, Sören Doose, and Markus Sauer for access to their TCSPC spectrometer, Suren Felekyan for providing the PDA software, and Johanna Rahm for coding support for the PDA analysis. We acknowledge funding by the CRC1507: Membrane-associated Protein Assemblies, Machineries, and Supercomplexes (Deutsche Forschungsgemeinschaft). H.H.N. acknowledges funding by the Deutsche Forschungsgemeinschaft (grant NI 694/3-1). S.M.A. and R.C. acknowledge the support of the Frankfurt Institute of Advanced Studies, the LOEWE Center for Multiscale Modelling in Life Sciences of the state of Hesse, and the International Max Planck Research School on Cellular Biophysics. R.C. acknowledges the support by Bayreuth University. S.M.A. and R.C. acknowledge computational resources and support by the Center for Scientific Computing of the Goethe University, and the Gauss Centre for Supercomputing e.V. (www.gauss-centre.eu) for funding this project by providing computing time on the GCS Supercomputer JUWELS at Jülich Supercomputing Centre (JSC).

## References

1. Lemmon, M. A., Schlessinger, J. & Ferguson, K. M. The EGFR family: not so prototypical receptor tyrosine kinases. Cold Spring Harb. Perspect. Biol. 6, a020768 (2014).

2. Yuzawa, S. et al. Structural basis for activation of the receptor tyrosine kinase KIT by stem cell factor. Cell 130, 323–334 (2007).

3. Opatowsky, Y. et al. Structure, domain organization, and different conformational states of stem cell factor-induced intact KIT dimers. Proc. Natl. Acad. Sci. U. S. A. 111, 1772–1777 (2014).

4. Chen, P.-H., Unger, V. & He, X. Structure of Full-Length Human PDGFRβ Bound to Its Activating Ligand PDGF-B as Determined by Negative-Stain Electron Microscopy. J. Mol. Biol. 427, 3921–3934 (2015).

5. Ognjenović, J., Grisshammer, R. & Subramaniam, S. Frontiers in Cryo Electron Microscopy of Complex Macromolecular Assemblies. Annu. Rev. Biomed. Eng. 21, 395–415 (2019).

6. Chung, I. et al. Spatial control of EGF receptor activation by reversible dimerization on living cells. Nature 464, 783–787 (2010).

7. Nagy, P., Claus, J., Jovin, T. M. & Arndt-Jovin, D. J. Distribution of resting and ligand-bound ErbB1 and ErbB2 receptor tyrosine kinases in living cells using number and brightness analysis. Proc. Natl. Acad. Sci. U. S. A. 107, 16524–16529 (2010).

8. Endres, N. F. et al. Conformational coupling across the plasma membrane in activation of the EGF receptor. Cell 152, 543–556 (2013).

9. Karathanasis, C. et al. Single-molecule imaging reveals the oligomeric state of functional TNFα-induced plasma membrane TNFR1 clusters in cells. Sci. Signal. 13, (2020).

10. Birchmeier, C., Birchmeier, W., Gherardi, E. & Vande Woude, G. F. Met, metastasis, motility and more. Nat. Rev. Mol. Cell Biol. 4, 915–925 (2003).

11. Trusolino, L., Bertotti, A. & Comoglio, P. M. MET signalling: principles and functions in development, organ regeneration and cancer. Nat. Rev. Mol. Cell Biol. 11, 834–848 (2010).

12. Organ, S. L. & Tsao, M.-S. An overview of the c-MET signaling pathway. Ther. Adv. Med. Oncol. 3, S7–S19 (2011).

13. Mellado-Gil, J. et al. Disruption of hepatocyte growth factor/c-Met signaling enhances pancreatic beta-cell death and accelerates the onset of diabetes. Diabetes 60, 525–536 (2011).

14. Campbell, D. B. et al. A genetic variant that disrupts MET transcription is associated with autism. Proc. Natl. Acad. Sci. U. S. A. 103, 16834–16839 (2006).

15. Shen, Y., Naujokas, M., Park, M. & Ireton, K. InIB-dependent internalization of Listeria is mediated by the Met receptor tyrosine kinase. Cell 103, 501–510 (2000).

16. Gherardi, E. et al. Functional map and domain structure of MET, the product of the c-met protooncogene and receptor for hepatocyte growth factor/scatter factor. Proc. Natl. Acad. Sci. U. S. A. 100, 12039–12044 (2003).

17. Uchikawa, E., Chen, Z., Xiao, G.-Y., Zhang, X. & Bai, X.-C. Structural basis of the activation of c-MET receptor. Nat. Commun. 12, 4074 (2021).

18. Cioce, V. et al. Hepatocyte growth factor (HGF)/NK1 is a naturally occurring HGF/scatter factor variant with partial agonist/antagonist activity. J. Biol. Chem. 271, 13110–13115 (1996).

19. Lemmon, M. A. & Schlessinger, J. Cell signaling by receptor tyrosine kinases. Cell 141, 1117–1134 (2010).

20. Jumper, J. et al. Highly accurate protein structure prediction with AlphaFold. Nature 596, 583–589 (2021).

21. Baek, M. et al. Accurate prediction of protein structures and interactions using a three-track neural network. Science 373, 871–876 (2021).

22. Altintas, D. M. et al. The PSI Domain of the MET Oncogene Encodes a Functional Disulfide Isomerase Essential for the Maturation of the Receptor Precursor. Int. J. Mol. Sci. 23, (2022).

23. Niemann, H. H. et al. Structure of the human receptor tyrosine kinase met in complex with the Listeria invasion protein InlB. Cell 130, 235–246 (2007).

24. Stamos, J., Lazarus, R. A., Yao, X., Kirchhofer, D. & Wiesmann, C. Crystal structure of the HGF beta-chain in complex with the Sema domain of the Met receptor. EMBO J. 23, 2325–2335 (2004).

25. Banerjee, M. et al. GW domains of the Listeria monocytogenes invasion protein InlB are required for potentiation of Met activation. Mol. Microbiol. 52, 257–271 (2004).

26. Souza, P. C. T. et al. Martini 3: a general purpose force field for coarse-grained molecular dynamics. Nat. Methods 18, 382–388 (2021).

27. Dietz, M. S., Wehrheim, S. S., Harwardt, M.-L. I. E., Niemann, H. H. & Heilemann, M. Competitive Binding Study Revealing the Influence of Fluorophore Labels on Biomolecular Interactions. Nano Lett. 19, 8245–8249 (2019).

28. Kalinin, S. et al. A toolkit and benchmark study for FRET-restrained high-precision structural modeling. Nat. Methods 9, 1218–1225 (2012).

29. Heilemann, M. et al. Subdiffraction-resolution fluorescence imaging with conventional fluorescent probes. Angew. Chem. Int. Ed Engl. 47, 6172–6176 (2008).

30. Petrelli, A. et al. The endophilin–CIN85–Cbl complex mediates ligand-dependent downregulation of c-Met. Nature 416, 187–190 (2002).

31. Baldering, T. N. et al. CRISPR/Cas12a-mediated labeling of MET receptor enables quantitative single-molecule imaging of endogenous protein organization and dynamics. iScience 24, 101895 (2021).

32. Harwardt, M.-L. I. E. et al. Single-Molecule Super-Resolution Microscopy Reveals Heteromeric Complexes of MET and EGFR upon Ligand Activation. Int. J. Mol. Sci. 21, (2020).

33. Harwardt, M.-L. I. E. et al. Membrane dynamics of resting and internalin B-bound MET receptor tyrosine kinase studied by single-molecule tracking. FEBS Open Bio 7, 1422–1440 (2017).

34. Petrelli, A. et al. Ab-induced ectodomain shedding mediates hepatocyte growth factor receptor down-regulation and hampers biological activity. Proc. Natl. Acad. Sci. U. S. A. 103, 5090–5095 (2006).

35. Duclos, C. M. et al. Caspase-mediated proteolysis of the sorting nexin 2 disrupts retromer assembly and potentiates Met/hepatocyte growth factor receptor signaling. Cell Death Discovery 3, 1–12 (2017).

36. Miekus, K. et al. MET receptor is a potential therapeutic target in high grade cervical cancer. Oncotarget 6, 10086–10101 (2015).

37. Dietz, M. S. et al. Single-molecule photobleaching reveals increased MET receptor dimerization upon ligand binding in intact cells. BMC Biophys. 6, 6 (2013).

38. Kapanidis, A. N. et al. Fluorescence-aided molecule sorting: analysis of structure and interactions by alternating-laser excitation of single molecules. Proc. Natl. Acad. Sci. U. S. A. 101, 8936–8941 (2004).

39. Preus, S., Noer, S. L., Hildebrandt, L. L., Gudnason, D. & Birkedal, V. iSMS: single-molecule FRET microscopy software. Nat. Methods 12, 593–594 (2015).

40. Lee, N. K. et al. Accurate FRET measurements within single diffusing biomolecules using alternating-laser excitation. Biophys. J. 88, 2939–2953 (2005).

41. Antonik, M., Felekyan, S., Gaiduk, A. & Seidel, C. A. M. Separating structural heterogeneities from stochastic variations in fluorescence resonance energy transfer distributions via photon distribution analysis. J. Phys. Chem. B 110, 6970–6978 (2006).

42. Kalinin, S., Sisamakis, E., Magennis, S. W., Felekyan, S. & Seidel, C. A. M. On the origin of broadening of single-molecule FRET efficiency distributions beyond shot noise limits. J. Phys. Chem. B 114, 6197–6206 (2010).

43. Montepietra, D., et al. FRETpredict: A Python package for FRET efficiency predictions using rotamer libraries. bioRxiv (2023) doi:10.1101/2023.01.27.525885.

44. Sasaki, T. et al. Structural basis for Gas6-Axl signalling. EMBO J. 25, 80–87 (2006).

45. Mohammadi, M., Olsen, S. K. & Ibrahimi, O. A. Structural basis for fibroblast growth factor receptor activation. Cytokine Growth Factor Rev. 16, 107–137 (2005).

46. Krimmer, S. G. et al. Cryo-EM analyses of KIT and oncogenic mutants reveal structural oncogenic plasticity and a target for therapeutic intervention. Proc. Natl. Acad. Sci. U. S. A. 120, e2300054120 (2023).

47. Niemann, H. H., Gherardi, E., Bleymüller, W. M. & Heinz, D. W. Engineered variants of InlB with an additional leucine-rich repeat discriminate between physiologically relevant and packing contacts in crystal structures of the InlB:MET complex. Protein Sci. 21, 1528–1539 (2012).

48. Niemann, H. H. et al. X-ray and Neutron Small-Angle Scattering Analysis of the Complex Formed by the Met Receptor and the Listeria monocytogenes Invasion Protein InlB. J. Mol. Biol. 377, 489–500 (2008).

49. Ferraris, D. M., Gherardi, E., Di, Y., Heinz, D. W. & Niemann, H. H. Ligand-mediated dimerization of the Met receptor tyrosine kinase by the bacterial invasion protein InlB. J. Mol. Biol. 395, 522–532 (2010).

50. Jaumouillé, V. & Waterman, C. M. Physical Constraints and Forces Involved in Phagocytosis. Front. Immunol. 11, 1097 (2020).

51. Niemann, H. H. Structural basis of MET receptor dimerization by the bacterial invasion protein InlB and the HGF/SF splice variant NK1. Biochimica et Biophysica Acta (BBA) - Proteins and Proteomics 1834, 2195–2204 (2013).

52. Andres, F. et al. Inhibition of the MET Kinase Activity and Cell Growth in MET-Addicted Cancer Cells by Bi-Paratopic Linking. J. Mol. Biol. 431, 2020–2039 (2019).

53. Hellenkamp, B. et al. Precision and accuracy of single-molecule FRET measurements-a multi-laboratory benchmark study. Nat. Methods 15, 669–676 (2018).

54. Agam, G. et al. Reliability and accuracy of single-molecule FRET studies for characterization of structural dynamics and distances in proteins. Nat. Methods 20, 523–535 (2023).

55. Schnitzbauer, J., Strauss, M. T., Schlichthaerle, T., Schueder, F. & Jungmann, R. Super-resolution microscopy with DNA-PAINT. Nat. Protoc. 12, 1198–1228 (2017).

56. Endesfelder, U., Malkusch, S., Fricke, F. & Heilemann, M. A simple method to estimate the average localization precision of a single-molecule localization microscopy experiment. Histochem. Cell Biol. 141, 629–638 (2014).

57. Ester, M., Kriegel, H. P., Sander, J. & X, Xu. A density-based algorithm for discovering clusters in large spatial databases with noise. in Proceedings of 2nd International Conference on Knowledge Discovery and Data Mining (KDD-96) (eds. Evangelos, S., Jiawei, H. & M., F. U.) 226–231 (AAAI Press, 1996).

58. Vogelsang, J. et al. A reducing and oxidizing system minimizes photobleaching and blinking of fluorescent dyes. Angew. Chem. Int. Ed Engl. 47, 5465–5469 (2008).

59. Sotolongo Bellón, J., et al. Four-color single-molecule imaging with engineered tags resolves the molecular architecture of signaling complexes in the plasma membrane. Cell Rep Methods 2, 100165 (2022).

60. Edelstein, A. D. et al. Advanced methods of microscope control using μManager software. J Biol Methods 1, (2014).

61. Cooper, M. et al. Cy3B: improving the performance of cyanine dyes. J. Fluoresc. 14, 145–150 (2004).

62. Lambert, T. J. FPbase: a community-editable fluorescent protein database. Nat. Methods 16, 277–278 (2019).

63. Muschielok, A. et al. A nano-positioning system for macromolecular structural analysis. Nat. Methods 5, 965–971 (2008).

64. Klose, D. et al. Resolving distance variations by single-molecule FRET and EPR spectroscopy using rotamer libraries. Biophys. J. 120, 4842–4858 (2021).

65. Schindelin, J., Rueden, C. T., Hiner, M. C. & Eliceiri, K. W. The ImageJ ecosystem: An open platform for biomedical image analysis. Mol. Reprod. Dev. 82, 518–529 (2015).

66. Eswar, N., Eramian, D., Webb, B., Shen, M.-Y. & Sali, A. Protein structure modeling with MODELLER. Methods Mol. Biol. 426, 145–159 (2008).

67. Pettersen, E. F. et al. UCSF Chimera--a visualization system for exploratory research and analysis. J. Comput. Chem. 25, 1605–1612 (2004).

68. Hu, X. et al. Structural and Functional Insight Into the Glycosylation Impact Upon the HGF/c-Met Signaling Pathway. Front Cell Dev Biol 8, 490 (2020).

69. Van Der Spoel, D. et al. GROMACS: fast, flexible, and free. J. Comput. Chem. 26, 1701–1718 (2005).

70. Vanommeslaeghe, K. et al. CHARMM general force field: A force field for drug-like molecules compatible with the CHARMM all-atom additive biological force fields. J. Comput. Chem. 31, 671–690 (2010).

71. Huang, J. & MacKerell, A. D., Jr. CHARMM36 all-atom additive protein force field: validation based on comparison to NMR data. J. Comput. Chem. 34, 2135–2145 (2013).

72. Varadi, M. & Velankar, S. The impact of AlphaFold Protein Structure Database on the fields of life sciences. Proteomics e2200128 (2022).

73. Brooks, B. R. et al. CHARMM: the biomolecular simulation program. J. Comput. Chem. 30, 1545–1614 (2009).

74. Michaud-Agrawal, N., Denning, E. J., Woolf, T. B. & Beckstein, O. MDAnalysis: a toolkit for the analysis of molecular dynamics simulations. J. Comput. Chem. 32, 2319–2327 (2011).

75. Wassenaar, T. A., Ingólfsson, H. I., Böckmann, R. A., Peter Tieleman, D. & Marrink, S. J. Computational Lipidomics with insane: A Versatile Tool for Generating Custom Membranes for Molecular Simulations. (2015) doi:10.1021/acs.jctc.5b00209.

76. van der Walt, S., Colbert, S. C. & Varoquaux, G. The NumPy array: A structure for efficient numerical computation. Comput. Sci. Eng. 13, 22–30 (2011).

77. Berman, H., Henrick, K., Nakamura, H. & Markley, J. L. The worldwide Protein Data Bank (wwPDB): ensuring a single, uniform archive of PDB data. Nucleic Acids Res. 35, D301–3 (2007).

78. Shaw, R. A., Johnston-Wood, T., Ambrose, B., Craggs, T. D. & Hill, J. G. CHARMM-DYES: Parameterization of Fluorescent Dyes for Use with the CHARMM Force Field. J. Chem. Theory Comput. 16, 7817–7824 (2020).

