## Supplementary Material for "Single-molecule imaging and molecular dynamics simulations reveal early activation of the MET receptor *in situ*"

Short Title: *In situ* early activation steps of MET receptor

Yunqing Li*[,1], Serena Arghittu*[,2,3], Marina S. Dietz[1], Gabriel J. Hella[2], Daniel Haße,[4] Davide M. Ferraris,[5] Petra Freund[1], Hans-Dieter Barth[1], Hartmut H. Niemann[4], Roberto Covino[2,3,6,#], Mike Heilemann[1,3,#]

*These authors equally contributed to this work.

### This PDF file includes:

Supplementary Notes 1 and 2
Figures S1 to S9
Tables S1 to S2

**Supplementary Note 1. MET dimerization, internalization, and detection probability in smFRET experiments**

Binding of InlB to MET receptors induces the formation of a dimeric complex $(InlB:MET)_2$. MET dimers were previously observed in cells using various microscopy methods, e.g., single-molecule photobleaching[1], quantitative single-molecule localization microscopy (qSMLM)[2], FRET-FLIM and fluorescence correlation spectroscopy (FCS)[3].

Upon ligand stimulation, receptor internalization reduces the membrane density of MET. A previous study showed that stimulation of human mammary epithelial cells (T47D/MET) with 100 ng/mL HGF or InlB for 15 min at 37°C (as used in this study) reduced the concentration of MET on the cell membrane by approximately 50%[4].

In an smFRET experiment, further considerations are necessary to estimate the theoretical detection probability. First, we used a total concentration of 10 nM InlB, equal amounts (5 nM) of donor- and acceptor-labeled InlB. The dissociation constant of fluorophore-labeled InlB from a InlB:MET complex is approximately 5 nM[1], which translates into a labeling efficiency of MET on the plasma membrane of approximately 65%. (Note that this assumes that the complex stability *in situ* is similar to that determined *in vitro*.) Second, the probability of donor/acceptor and acceptor/donor labeled $(MET:InlB)_2$ dimers is 25% each, or 50% in sum; only those complexes are accessible for smFRET measurements. Third, the degree of labeling of InlB affects the detectability. While we obtained stoichiometric labeling for ATTO 647N-H-$InlB_{321}$, other InlB constructs had a slightly lower degree of labeling (70-80%) (see **Supplementary Table 1**). Fourth, the total amount of donor/acceptor labeled $(MET:InlB)_2$ dimers might be reduced by incomplete dimerization (albeit that might be small; see Baldering et al.[2]), and by internalization of $(MET:InlB)_2$ dimers (see above).

**Supplementary Note 2. Comparison of H-T and T-H FRET pairs**

smFRET experiments that targeted the H-T distance were conducted separately, using either Cy3B-H-$InlB_{321}$ and ATTO 647N-T-$InlB_{321}$ or Cy3B-T-$InlB_{321}$ and ATTO 647N-H-$InlB_{321}$. From the E,S-histograms of H-T and T-H, FRET efficiencies of 0.594 ± 0.009 and 0.525 ± 0.005 were determined, respectively (**Supplementary Fig. S4**). We explain the difference in FRET efficiencies by a different rotational flexibility (and thus accessible volume) for the donor fluorophore in the H and T position. In the H-T pair, the donor dye is labeled at the H position of $InlB_{321}$, which faces to the outside of the protein complex (**Figure 2A**). This position provides more rotational freedom to the fluorophore, leading to a broader FRET distribution relative to the T-H combination and a marginally higher FRET efficiency (**Supplementary Fig. S4**). In addition, due to the slight asymmetry of the $(InlB:MET)_2$ dimer, the two H-T distances are slightly different (**Figure 2A**); this leads to an additional broadening of the FRET distribution, as compared to the T-T distance (**Figure 4A**). The data sets for H-T and T-H were merged, to average out the bias of two possible configurations.

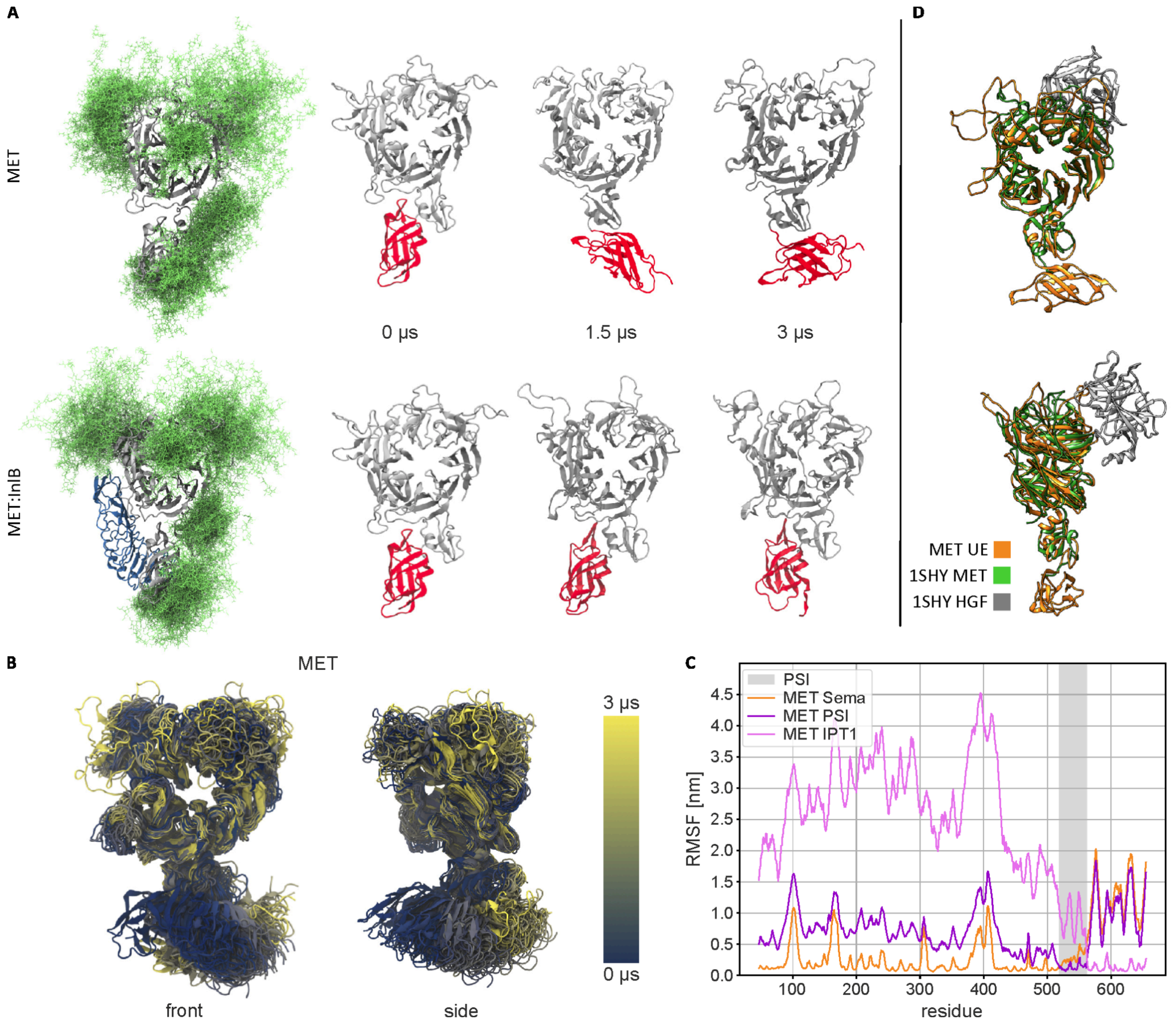


**Supplementary Figure 1.** (A) Renders of the N-glycosylated MET upper ectodomain system in isolation (top row, MET upper ectodomain) and bound to InlB (bottom row, MET upper ectodomain:InlB). The renders show the MD models of the MET receptor and three snapshots of the corresponding trajectories. In the first column, the models of the glycosylated MET in isolation and glycosylated MET in complex with InlB are shown (InlB in blue, MET in silver, and in green glycan conformations from the first 200 ns of the trajectory, sampled every 2.5 ns; water and ions are not shown for clarity). In the other columns, only the MET receptor is shown for both models at different time points. Sema and PSI are represented in silver, the IPT1 domain is in red. Glycans, water, and ions are not shown for clarity. (B) Side and front view of the MET upper ectodomain along the model trajectory (every 50 frames). The cartoon representation is colored according to the frame index (according to the color bar). (C) Flexibility of the MET upper ectodomain in relation to its various domains. The root mean square fluctuation (RMSF) of the Cα atom positions within the MET upper ectodomain with respect to the initial frame. The trajectory was aligned to the Sema (orange), PSI (dark purple), and IPT1 (light purple) domain, respectively. The PSI domain residues are highlighted by the gray area.

(D) Alignment of the MET:HGF structure (PDB 1SHY, MET in green, HGF in silver) to the MET upper ectodomain in the last frame of the simulation (MET in orange).

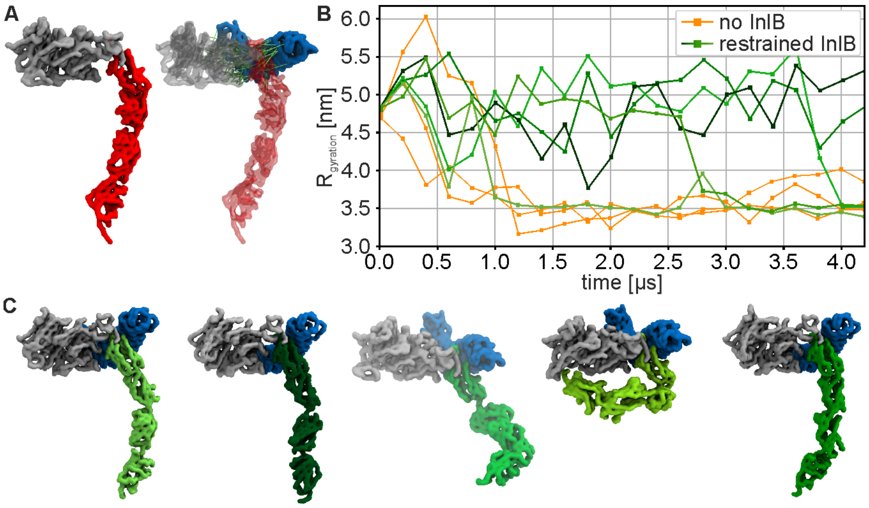


**Supplementary Figure 2. Coarse-grain study of the conformational plasticity of the entire MET ectodomain.** (A) Renders of the coarse-grained (CG) models of MET (left) and of the restrained MET:InlB$_{321}$ complex (right). The receptor is represented as a surface, the SEMA and PSI are in silver and the MET's stalk in red. In the restrained complex the receptor is transparent, InlB is in blue and the restraints are in lime green (the restraints are shown 1 every 20). (B) Radius of gyration of the MET (orange) and of the restrained MET:InlB$_{321}$ complex (green shades) replicas. Same as in Fig. 1I and reported here for the convenience of the reader. (C) Snapshots from the 5 replica trajectories of the restrained MET:InlB$_{321}$ complex showing different modes of the receptor's stalk. The snapshots are taken at 2.5 $\mu s$ The complex is represented as a surface; SEMA, PSI are in silver, InlB in blue and the MET's stalk is in different shades of green corresponding to the different replicas.

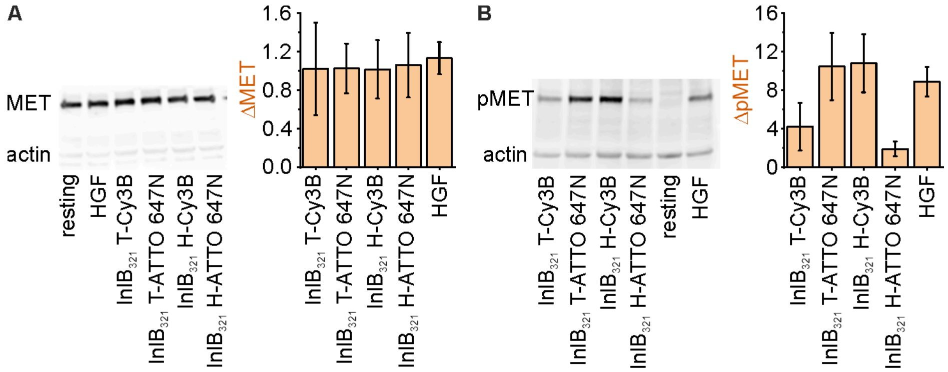


**Supplementary Figure 3. Western blot analysis of MET in resting and ligand-stimulated U-2 OS cells.** Exemplary western blots of (A) MET and (B) phosphorylated MET (pMET) are shown (left). Cells were incubated for 15 min with ligand ($InlB_{321}$ or HGF) or medium only for the resting condition. Actin was co-labeled for quantification. Page ruler was used as a size marker. In the bar graphs (right) the relative difference in the amount of MET and pMET, respectively, is shown. For this purpose, the MET bands were normalized with reference to the actin bands. The ligand-activated cells were compared to the resting cells. 3-4 independent experiments were averaged. Errors are given as standard deviations.

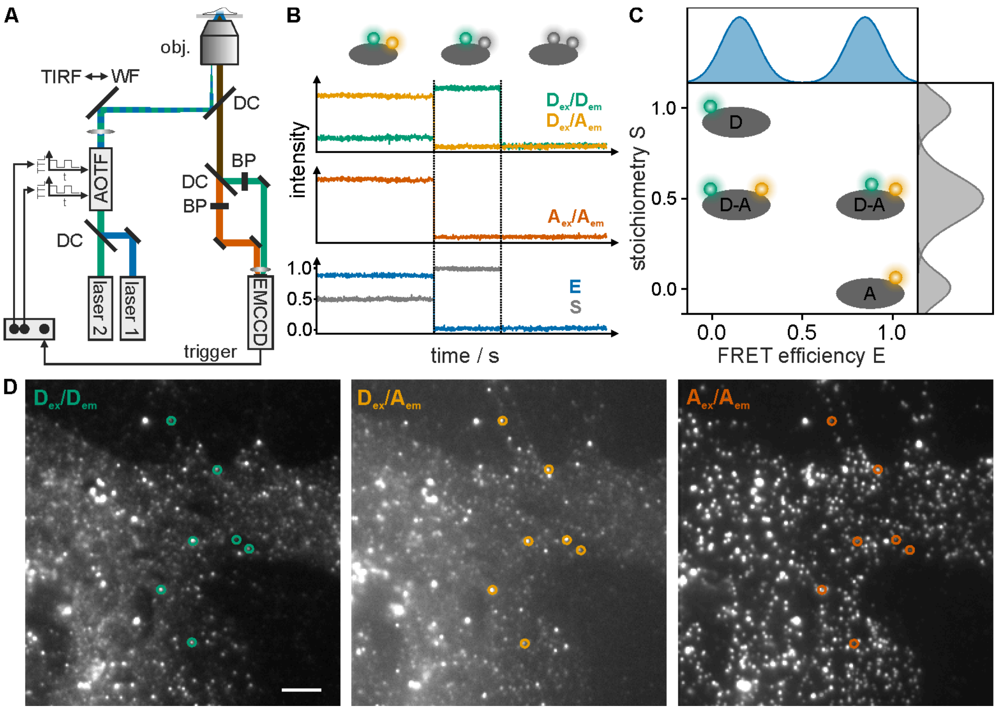


**Supplementary Figure 4. Single-molecule FRET with alternating laser excitation (ALEX).** (A) Scheme of a microscope setup for single-molecule FRET measurements with alternating laser excitation. A donor and an acceptor excitation laser are alternated using an acousto-optical filter (AOTF). An adjustable mirror is used to adjust illumination to total internal reflection (TIRF) to solely illuminate the lower plasma membrane of the cells and reduce background fluorescence. DC: dichroic mirror, obj.: objective, BP: bandpass filter. (B) Schematic single FRET pair traces. Shown are the donor (green) and acceptor emission (light orange) upon donor excitation and the acceptor emission (dark orange) upon acceptor excitation. From these intensities the FRET efficiency E (blue) and the stoichiometry S (gray) of donor and acceptor can be calculated. On the vertical lines, the donor or acceptor photobleach, which can be seen in the intensity traces as well as the E and S traces. (C) Two-dimensional ALEX histogram showing the expected populations for different cases. In the case of active donor and acceptor, a stoichiometry of 0.5 is expected and the FRET efficiency relates to the distance between donor and acceptor. In a scenario where only the donor is present; a stoichiometry of 1 is expected, while a molecule having only an active acceptor exhibits a stoichiometry of 0. (D) Exemplary smFRET data of $InlB_{321}$-labeled MET receptors in U-2 OS cells. $D_{ex}/D_{em}$ shows donor emission upon donor excitation. $D_{ex}/A_{em}$ shows acceptor emission upon donor excitation. $A_{ex}/A_{em}$ shows acceptor emission upon acceptor excitation. The image is the average intensity of the first 100 frames. FRET pairs are highlighted by circles. Scale bar 5 µm.

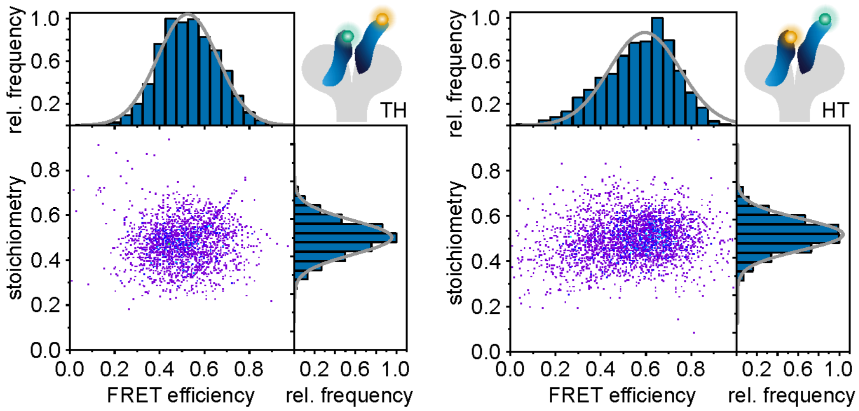


**Supplementary Figure 5. Single-molecule E,S-histogram obtained from T-H/H-T smFRET experiments of $(MET:InlB)_2$ dimers in U-2 OS cells.** Left: E,S-histogram for Cy3B-T-$InlB_{321}$ and ATTO 647N-H-$InlB_{321}$ (N = 25 smFRET traces from 20 cells); right: E,S-histogram for Cy3B-H-$InlB_{321}$ and ATTO 647N-T-$InlB_{321}$ (N = 24 smFRET traces from 19 cells).

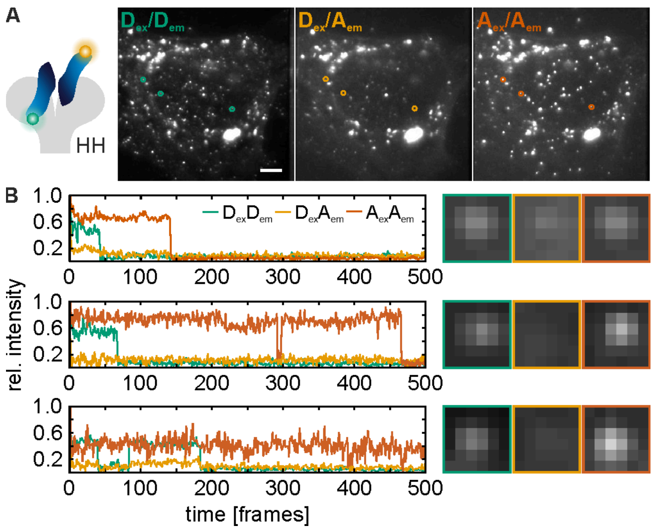


**Supplementary Figure 6. Exemplary intensity traces for InlB H-Cy3B and H-ATTO 647N variants.** (A) Exemplary smFRET data of $InlB_{321}$-labeled MET receptors in U-2 OS cells. $D_{ex}/D_{em}$ shows donor emission upon donor excitation (green). $D_{ex}/A_{em}$ shows acceptor emission upon donor excitation (light orange). $A_{ex}/A_{em}$ shows acceptor emission upon acceptor excitation (dark orange). The images are the average intensities of the first 100 frames. The donor-acceptor pairs shown in (B) are highlighted by circles. Scale bar 5 µm. (B) The intensity traces for the donor and the acceptor upon donor excitation as well as the acceptor intensity upon acceptor excitation are shown. No FRET signal is observed in the $D_{ex}A_{em}$ channel. Traces are normalized to 1.

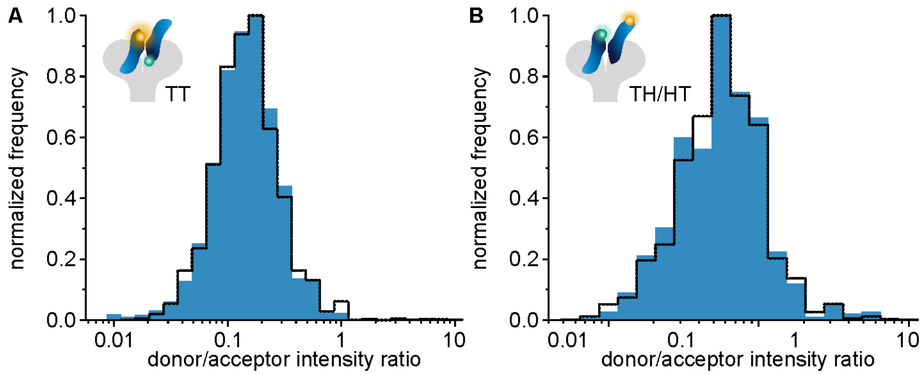


**Supplementary Figure 7. Photon distribution analysis (PDA) of T-T (A) and H-T/T-H FRET measurements (B).** In blue the histograms of the ratio of donor emission and acceptor emission upon donor excitation are shown. The intensities are not background corrected. The dashed line represents the simulated histograms according to Antonik et al.[5] with a single-state model.

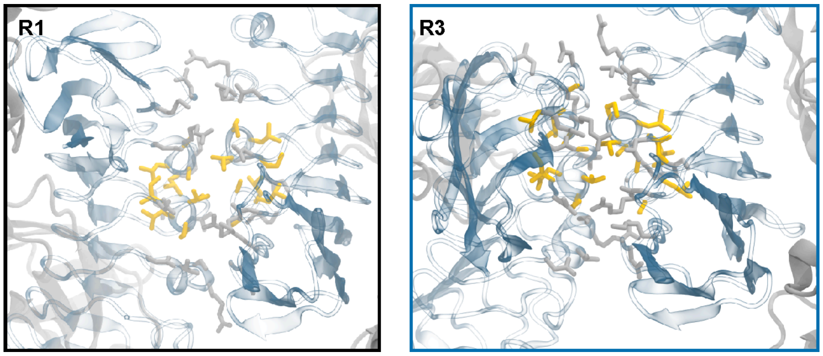


**Supplementary Figure 8. Hydrophobic core of the dimer interaction interface in replica 1 and 3.** Hydrophobic amino acids are highlighted in yellow. InlB and MET are shown as a transparent cartoon, respectively blue and gray. The residues involved in the interfaces are shown in licorice representation, hydrophobic residues are colored in yellow and non-hydrophobic in silver.

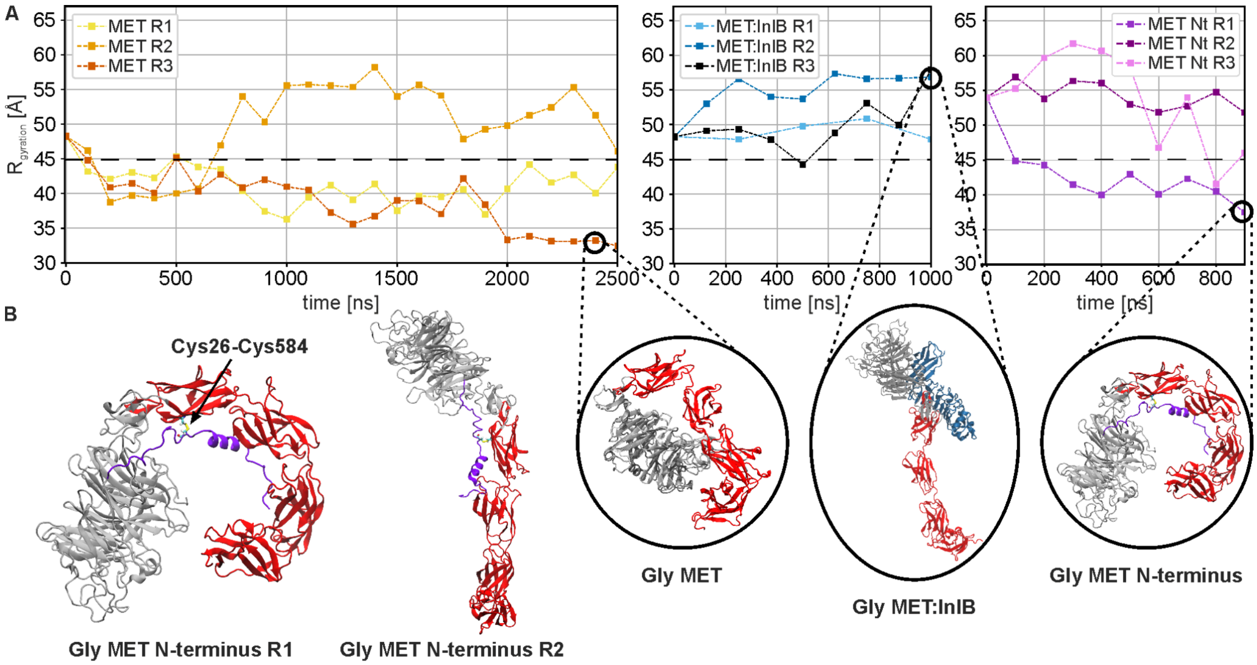


**Supplementary Figure 9.** (A) Radius of gyration of, from left to right, the isolated MET receptor ectodomain, the MET:InlB$_{321}$ complex, the isolated MET receptor ectodomain with an additional 42 residue N-terminal loop and a disulfide bond between CYS26 and CYS584 on IPT1. In the inserts exemplary snapshots of each model from the end of a replica trajectory. First and second panels are the same as in Fig. 1H, and reported here for the convenience of the reader. (B) Exemplary renders showing the compact and extended conformations of the isolated MET with N-terminal loop and the CYS26-CYS584 .

**Supplementary Table 1. Used $InlB_{321}$ variants and the respective mutations for fluorophore labeling.** The degrees of labeling (DOL) of Cy3B- or ATTO 647N-labeled variants were determined by absorption spectroscopy.

| **Variant** | **Mutation** | **DOL (%)** |
|---|---|---|
| InlB-T-Cy3B | K280C | 87 |
| InlB-T-ATTO 647N | K280C | 69 |
| InlB-H-Cy3B | K64C | 70 |
| InlB-H-ATTO 647N | K64C | 103 |

**Supplementary Table 2. Density of MET receptor clusters in different cell lines.** Mean receptor densities were determined from *d*STORM super-resolution data and were corrected for background signals. The errors are standard deviations.

| **Cell line** | **Receptor density ($\mu m^2$)** |
|---|---|
| 23132/87 | 7.8 ± 2.5 |
| HeLa | 8.7 ± 2.7 |
| Huh7.5 | 3.5 ± 1.1 |
| U-2 OS | 2.8 ± 1.2 |
| U-251 | 10.6 ± 2.7 |

### References


1. Dietz, M. S. *et al.* Single-molecule photobleaching reveals increased MET receptor dimerization upon ligand binding in intact cells. *BMC Biophys.* **6**, 6 (2013).
2. Baldering, T. N. *et al.* CRISPR/Cas12a-mediated labeling of MET receptor enables quantitative single-molecule imaging of endogenous protein organization and dynamics. *iScience* **24**, 101895 (2021).
3. Koschut, D. *et al.* Live cell imaging shows hepatocyte growth factor-induced Met dimerization. *Biochim. Biophys. Acta* **1863**, 1552–1558 (2016).
4. Li, N., Hill, K. S. & Elferink, L. A. Analysis of receptor tyrosine kinase internalization using flow cytometry. *Methods Mol. Biol.* **457**, 305–317 (2008).
5. Antonik, M., Felekyan, S., Gaiduk, A. & Seidel, C. A. M. Separating structural heterogeneities from stochastic variations in fluorescence resonance energy transfer distributions via photon distribution analysis. *J. Phys. Chem. B* **110**, 6970–6978 (2006).
